# Data mining sheds light on a novel family of plant-associated negative-sense RNA viruses linked to lispiviruses

**DOI:** 10.64898/2026.09.18.752651

**Authors:** Nicolás Bejerman, David Huebert, Robert Alvarez-Quinto, Diego Quito-Avila, Humberto Debat

**Author notes:** Corresponding authors: NB,; HD. These authors contributed equally to this work (Nicolás Bejerman, David Huebert).

## Abstract

Large-scale mining of public transcriptomic datasets can reveal viral diversity that remains invisible to conventional virus-surveillance approaches, while increasingly powerful structure-prediction methods provide a complementary route to characterizing highly divergent viral proteins. Here, we combine sequence detection, phylogenetic analysis, structural prediction, and host-association analyses to investigate the cryptic diversity and biology of a new clade of plant-associated lispi-like viruses. Analyses of RNA-sequencing datasets identified and enabled assembly of 87 lispi-like virus genomes associated with 75 plant hosts, expanding the known diversity of this new group by approximately 40-fold. The viruses share a conserved four-cistron genome organization, 3′-N-P2-P3-L-5′. Structural analyses provide functional insights into the four conserved proteins. P1 (N) adopts a canonical negative-strand RNA virus nucleocapsid architecture with conserved RNA-interacting residues and a predicted RNA-packaging configuration. P2 is exceptionally divergent; although a subset of structures resembles the ITPase/HAM1 fold. P3 forms a conserved trimeric coiled-coil architecture reminiscent of a viral fusion-protein stalk, but lacks the family-wide sequence features expected of a canonical membrane glycoprotein. P4 contains a structurally resolved Mononegavirales-type RNA-dependent-RNA-polymerase (RdRp) core with invariant catalytic motifs, including the characteristic GDN signature, whereas its accessory regions are substantially more divergent. Phylogenetic insights form a distinct monophyletic lineage sister to the predominantly invertebrate-associated *Lispiviridae*, supporting their recognition as the new proposed family Masuviridae, comprising 15 tentative genera. Genus-level clustering is accompanied by marked differences in host association, ranging from strong specialization to broader host ranges. Retrospective screening of public sequencing libraries further identified lispi-like virus sequences in 1,536 libraries representing 134 plant species, 11 plant families and 182 geographic locations, highlighting a substantial and geographically widespread cryptic virome. Together, these results establish Masuviridae as a deeply divergent lineage of plant-associated negative-sense RNA viruses and illustrate how the integration of sequence, structural, and large-scale transcriptomic analyses can move viral dark matter from detection towards evolutionary and functional characterization.

## 1. Introduction

In the current era of metagenomics, the rapid discovery of novel viruses has revealed a vast and diverse evolutionary landscape of replicating entities, presenting significant challenges in their systematic classification [1]. In response to this complexity, various strategies have emerged, culminating in a comprehensive proposal for establishing a "megataxonomy" of the viral world [2].

Despite extensive efforts to characterize the viral component of the biosphere, it is evident that only a tiny fraction, likely less than one percent, of viral diversity has been thoroughly explored [3–6]. As a result, our understanding of the global virome remains limited, particularly regarding its immense diversity and the interactions between viruses and their hosts [7–10]. To address this gap, researchers have increasingly turned to mining publicly available transcriptome datasets derived from High-Throughput Sequencing (HTS), which offers a rapid and cost-effective approach to viral discovery [8, 11–14]. This data-driven strategy has proven particularly valuable given the sheer volume of freely available datasets within the Sequence Read Archive (SRA), maintained by the National Center for Biotechnology Information (NCBI), which continues to expand at an extraordinary pace. Although the data in SRA represent a partial and potentially biased sample of Earth’s biodiversity, the NCBI-SRA remains an exceptionally efficient and economical resource for the identification of novel viruses [15]. The tool ‘Serratus’ [8] has proven invaluable, enabling large-scale data mining and accelerating viral sequence discovery at an unprecedented pace.

In the realm of virus taxonomy, a growing consensus has emphasized the necessity of incorporating viruses identified solely through metagenomic data into the official classification framework of the International Committee on Taxonomy of Viruses (ICTV) [16]. This shift underscores the vital role of metagenomic approaches in expanding our understanding of the global virome and modernizing taxonomic systems to accommodate the ever-growing known diversity of viruses [17].

Negative-sense RNA viruses constitute a comparatively underexplored component of the plant virosphere [18]. In contrast to positive-sense RNA viruses, whose diversity has been extensively documented through both disease surveillance and metatranscriptomic surveys, plant-associated negative-sense viruses have historically been represented by a smaller number of taxonomic lineages [5-6, 18, https://ictv.global/taxonomy]. Recent transcriptome-mining studies, however, have begun to reveal a broader and more complex diversity of plant-associated negative-sense viruses, including viruses detected in asymptomatic hosts and lineages that lack established taxonomic or biological frameworks [11, 19–20]. This expanding diversity raises a fundamental question: to what extent does the apparent paucity of plant-associated negative-sense RNA virus lineages reflect biological rarity, and to what extent does it reflect limitations in sampling and sequence-based discovery?

The rapid expansion of viral sequence space has also exposed a second challenge: identifying a viral sequence is increasingly easier than determining what its encoded proteins do. Viral proteins often evolve rapidly, and sequence divergence can erase recognizable homology even when structural and functional constraints are retained [21–22]. This problem is particularly acute for newly discovered RNA viruses, for which many proteins have no reliable sequence-based annotation and therefore remain part of the viral functional “dark matter” [23–24]. Recent large-scale structural analyses have shown that predicted protein structures can recover deeply conserved folds and reveal relationships that are undetectable from primary sequence alone, providing a complementary route to functional and evolutionary inference [21–24]. Structure-guided approaches are therefore especially valuable for newly discovered viral lineages in which conventional homology searches provide little information, allowing conserved architectural features to be distinguished from genuinely lineage-specific proteins while also defining the limits of functional inference.

The order *Mononegavirales* was created to accommodate viruses with linear, single-stranded, negative-sense RNA genomes [25]. Currently, this order is composed of 11 families, where the only family to date to harbor plant-associated viruses is the *Rhabdoviridae* [26]. *Lispiviridae* is a family encompassing viruses with negative-sense RNA genomes of 6.5–15.5 kb which have mainly been identified in arthropods [27]. Genomes of members of the family *Lispiviridae* commonly have five to six open reading frames (ORFs). Encoded proteins likely include a nucleocapsid (N), a phosphoprotein (P), a glycoprotein (G) and a large protein (L) including an RNA-directed-RNA-polymerase (RdRP) domain [27]. Currently, the family *Lispiviridae* includes 30 genera with 45 species (https://ictv.global/taxonomy). In 2018, a novel rod-shaped virus with a negative-sense single-stranded genome was identified in maize samples collected from Ecuador. The virus, tentatively named maize suscal virus (MSuV), had a 12 kb genome comprising four ORFs with the predicted L protein phylogenetically related to lispiviruses [28]. Later, the L gene of a novel lispi-like virus associated to the plant host *Disporopsis* spp. was identified by [29], who proposed that both viruses may belong to a new sister lineage of the family *Lispiviridae*, for which we propose the name Masuviridae within the order *Mononegavirales*.

Here, we combine large-scale mining of public RNA-sequencing datasets with phylogenomic, host-association and structure-guided analyses to characterize the diversity and biology of plant-associated lispi-like viruses. We identified and assembled 87 lispi-like virus coding sequences associated with 75 plant hosts and use comparative phylogenetic analyses to establish their evolutionary relationship with Lispiviridae and to develop a proposed taxonomic framework for this lineage. We then examine the predicted structures of all four major viral proteins to distinguish conserved viral architectures from lineage-specific and unresolved regions and to generate testable hypotheses about their functions. Lastly, we use large-scale retrospective screening of public RNA-sequencing libraries to assess the broader distribution, host range and apparent prevalence of these viruses. Together, these analyses move beyond sequence-based discovery to provide an integrated view of the evolutionary, structural and ecological diversity of a previously obscure group of plant-associated negative-sense RNA viruses.

## 2. Material and Methods

### 2.1 Identification of lispi-like virus sequences from public plant RNA-seq datasets

Plant-associated lispi-like virus sequences were identified by mining publicly available transcriptomic datasets. The Serratus database was searched using the Serratus Explorer [8], with the predicted RNA-dependent RNA polymerase (RdRp) protein of maize suscal virus (MSuV; NCBI RefSeq) used as the primary query. SRA libraries showing matches to the query sequence were retained using an alignment identity threshold >45% and a score >10. In parallel, the NCBI Transcriptome Shotgun Assembly (TSA) database was searched using the amino acid sequences of the MSuV nucleocapsid (N) and polymerase (L) proteins as queries in tBLASTn searches against Viridiplantae (taxid:33090). TSA searches used word size 6, an E-value threshold of 10, and the BLOSUM62 scoring matrix.

### 2.2. Sequence retrieval, assembly and genome reconstruction

Raw sequence reads from SRA experiments identified through Serratus were retrieved from the corresponding NCBI BioProjects (Tables 1–4). Reads were quality- and adapter-trimmed using Trimmomatic v0.40 [30], using standard parameters except that the quality threshold was increased from 20 to 30. The initial ILLUMINACLIP step and sliding-window quality trimming were applied, with a required average quality of 30. Filtered reads were assembled de novo using rnaSPAdes with standard parameters on the Galaxy server. Resulting transcripts were screened by local BLASTX searches against MSuV RefSeq proteins using an E-value threshold of 1 × 10^−5^. Candidate viral contigs were examined manually and extended and/or validated by iterative mapping of filtered reads from the corresponding SRA library. In each iteration, reads mapping to the candidate contig were extracted and used to extend the sequence, after which the extended sequence was used as the query for the subsequent iteration. Iterations were continued until no further extension was obtained. Final extended sequences were reassembled and polished using Geneious v8.1.9 (Biomatters Ltd.) with high-sensitivity alignment parameters. Sequences were retained for downstream analysis when a complete coding region could be reconstructed. The resulting dataset comprised 87 lispi-like virus coding sequences, including distinct variants associated with six viruses. Sequence completeness and coding-region organization were assessed by inspection of read-supported assemblies and predicted open reading frames.

### 2.3. Genome annotation and comparative sequence analysis

Open reading frames (ORFs) were predicted using NCBI ORFfinder, using genetic code 1 and a minimum ORF length of 120 nt. Predicted proteins were annotated using the InterPro protein sequence analysis platform and the NCBI Conserved Domain Database (CDD v3.20), using an E-value threshold of 0.01. Divergent proteins were further examined using HHpred and HHblits as implemented in the MPI Bioinformatics Toolkit. Homologous proteins were identified using BLASTP searches against the NCBI non-redundant (nr) protein database. Transmembrane topology was initially assessed using TMHMM v2.0. Signal peptides were predicted using SignalP 6.0 [31]. Where appropriate, combined signal-peptide and transmembrane-topology predictions were assessed using Phobius [32] Nuclear localization signals were predicted using cNLS Mapper. All web-based prediction tools were run using their standard parameters unless otherwise specified. Protein structures were additionally searched against publicly available structural databases using Foldseek [33].

### 2.4. Analysis of genome organization and gene junctions

The genomic organization of the 87 lispi-like virus sequences was determined from the reconstructed coding regions and comparative nucleotide alignments. Gene boundaries and the relative positions of the N, P2, P3, and L ORFs were compared across all sequences. To characterize conserved transcription-associated elements, nucleotide sequences surrounding each gene junction were aligned and inspected in the mRNA-sense orientation. Conserved 3′ termination/polyadenylation signals, intergenic spacers, and downstream transcription-initiation motifs were identified by comparative analysis of all available gene junctions. Motif frequencies and sequence conservation were calculated across the 87 sequences using standard sequence-alignment and nucleotide-counting procedures.

### 2.5. Pairwise amino acid sequence identity

Pairwise amino acid sequence identities were calculated for the predicted L proteins of the lispi-like viruses identified in this study and selected related viruses available in the NCBI database. Multiple sequence alignments were generated using MAFFT v7.505 [34] with standard parameters, and pairwise identities were calculated using SDTv1.2 [35]. For comparative analyses of individual viral proteins, the corresponding amino acid sequences were aligned using MAFFT with standard parameters. Mean, median and range of pairwise identities were calculated from the resulting alignments.

### 2.6. Phylogenetic analysis and taxonomic classification

Phylogenetic analyses were performed using the predicted L protein sequences of the 87 plant-associated lispi-like viruses together with MSuV, Disporopsis pernyi-associated lispi-like virus and selected representatives of the family Lispiviridae. Selected rhabdovirus L proteins were included as an outgroup. Amino acid sequences were aligned using MAFFT v7.505. The resulting alignments were analyzed using MEGA11 [36]. Maximum-likelihood phylogenetic trees were inferred using the WAG+G+F substitution model, and node support was estimated from 1,000 bootstrap replicates. Tree topology and branch support were visualized using standard phylogenetic visualization procedures. Provisional species and genus boundaries were evaluated using the combination of phylogenetic clustering and pairwise amino acid identity of the L protein. An L-protein amino acid identity threshold of 80% was used as a provisional species-level criterion within the newly proposed genera, based on the distribution of sequence identities observed among the sampled viruses. Genus-level groups were defined by discrete monophyletic clustering in the L-protein phylogeny together with marked inter-group sequence divergence.

### 2.7. Protein structure prediction

Three-dimensional structures of the four major viral proteins were predicted to investigate conserved structural features that could not be reliably inferred from primary sequence similarity. AlphaFold2 [37] was used to generate predicted structures for the P1, P2, and P3 proteins of the 87 lispi-like viruses. Structural predictions were generated using standard parameters. Prediction confidence was assessed using per-residue predicted local distance difference test (pLDDT) scores and, where available, predicted alignment error (PAE). Regions with low confidence were interpreted cautiously and were not considered evidence of a defined three-dimensional fold. For P3, oligomeric models were additionally generated using AlphaFold-Multimer [38] with standard parameters. Homotrimeric models were evaluated for consistency of the predicted coiled-coil architecture and overall assembly. A representative P4/L protein was modeled using AlphaFold2 and compared with experimentally determined structures of Mononegavirales polymerases. The representative sequence was selected to provide a structurally informative model while avoiding redundant structural analyses of highly similar variants.

### 2.8. Structure-based protein annotation

Predicted protein structures were searched against experimentally determined structures in the Protein Data Bank and predicted structures in the AlphaFold Protein Structure Database using Foldseek [33]. Searches were performed using standard parameters. Structural matches were evaluated according to the significance and extent of structural alignment, together with agreement between the predicted fold and known protein architecture. Structural assignments were considered putative when supported by a coherent fold-level match but were not interpreted as definitive functional annotations in the absence of independent sequence, catalytic or structural evidence. For P1, representative predicted structures were compared with experimentally determined nucleoprotein structures from negative-sense RNA viruses. Structural superposition was performed using standard structural-alignment procedures, and conserved residues within structurally corresponding RNA-interacting regions were identified from the multiple sequence alignment and structural models. For P4, a representative AlphaFold2 model was compared with the vesicular stomatitis virus L protein structure [PDB 5A22; 39], using structural superposition and Foldseek. The organization of the fingers, palm and thumb subdomains of the RdRp core was assessed, and the positions of the canonical polymerase motifs were examined relative to the experimentally determined Mononegavirales polymerase architecture.

### 2.9. Functional assessment of P2

Because conventional sequence searches failed to provide reliable functional annotation for P2, all 87 P2 structures were subjected to structure-based searches against the Protein Data Bank and AlphaFold Protein Structure Database using Foldseek. Putative functional assignments were classified according to the structural matches returned by Foldseek. Particular attention was given to structures related to the inosine triphosphate pyrophosphatase (ITPase)/Maf/HAM1 superfamily. The ITPase-like interpretation was independently evaluated using profile hidden Markov models. P2 sequences were searched against the Pfam Ham1 (PF01725) and Maf (PF02545) profiles using standard HMM-based search parameters. Human ITPA, Escherichia coli RdgB and Saccharomyces cerevisiae Ham1 proteins were included as positive controls for the corresponding nucleotide-pool sanitization proteins. The presence or absence of significant profile matches was compared between the control proteins and the lispi-like P2 sequences. Sequence conservation within the P2 family was assessed from a multiple sequence alignment generated using MAFFT with standard parameters. Conserved residues were identified from the resulting alignment. For ITPase-like structures, residues corresponding to conserved catalytic or substrate-binding positions in characterized ITPase/HAM1 proteins were mapped onto the predicted structures. In particular, the structural equivalents of the conserved catalytic lysine were examined. Potential substrate-binding pockets in ITPase-like P2 models were inspected qualitatively by comparison with characterized ITPase/HAM1 structures. No molecular docking or quantitative ligand-binding calculation was performed. Structural observations were therefore interpreted as hypotheses concerning possible substrate recognition rather than evidence of enzymatic activity.

### 2.10. Analysis of P3 topology and structural features

P3 proteins were examined as the third ORF in the conserved four-cistron genome organization. Predicted P3 structures were analyzed both as monomers and as homotrimers using AlphaFold2 and AlphaFold-Multimer, respectively. The predicted trimeric architecture was assessed for the presence of conserved helical regions, coiled-coil organization, and overall similarity to known viral fusion-protein architectures. Structural comparisons were performed using standard structural-alignment procedures. Potential membrane-associated features were assessed using Phobius, TMHMM and SignalP 6.0. N-terminal signal peptides and transmembrane helices were recorded for each sequence. Potential N-glycosylation sites were identified from canonical N-X-S/T sequons, where X is any amino acid other than proline. Potential proteolytic cleavage motifs were assessed by searching the aligned sequences for conserved sequence patterns associated with known plant secretory-pathway proteases, using standard motif-based searches. The distribution of these features was evaluated across the complete 87-sequence dataset and within individual phylogenetic/genus-level groups.

### 2.11. Selection and sequence-conservation analyses

Selection analyses of P3 were performed using codon-based alignments generated from the corresponding nucleotide and amino acid sequences. Protein alignments were used to guide the corresponding nucleotide alignment, and poorly aligned or ambiguous regions were excluded from downstream calculations. Synonymous and nonsynonymous substitutions were estimated using the Nei–Gojobori method [40]. Because synonymous substitutions were saturated across the full P3 dataset, dN/dS estimates were restricted to low-divergence sequence pairs for which synonymous divergence remained interpretable. Mean dN/dS values were calculated for the complete protein and for structurally defined regions, including the conserved coiled-coil and the more variable head/loop regions. Site-wise selection was assessed using within-cluster comparisons to reduce the effects of extreme sequence divergence and synonymous-site saturation. Codons were classified as showing evidence of purifying selection when the estimated synonymous/nonsynonymous substitution pattern met the statistical significance criterion used in the analysis; diversifying sites were identified using the corresponding criterion. Multiple-testing correction was applied where appropriate.

For P4, sequence conservation was quantified from the multiple sequence alignment using per-column amino acid entropy. Shannon entropy was calculated for each alignment position, with lower values indicating greater conservation. Conservation profiles were compared between the structurally resolved RdRp core and the more divergent C-terminal region using a one-sided Mann–Whitney U test.

### 2.12. Analysis of P4 polymerase motifs

The predicted P4/L proteins were examined for the canonical catalytic motifs of Mononegavirales RNA-dependent RNA polymerases. Conserved sequence blocks corresponding to palm motifs A, B and C were identified from the multiple sequence alignment and compared with experimentally characterized Mononegavirales polymerases. Particular attention was given to the catalytic residues of motif A and the motif-C GDN signature characteristic of non-segmented negative-sense RNA virus polymerases. The conservation of these residues and their structural positions within the predicted RdRp core were assessed across all 87 P4 sequences. The structural organization of the P4 protein was compared with the L protein structure of vesicular stomatitis virus (VSV, PDB 5A22), with particular attention to the fingers, palm and thumb subdomains and the relative position of the catalytic motifs.

### 2.13. Host-association and genus-level analyses

Host information associated with the viral sequences was obtained from the corresponding SRA/TSA metadata and source datasets. Host plant names were standardized to current taxonomic nomenclature and assigned to their corresponding botanical families. Virus–host associations were summarized at the species and family levels for each proposed viral genus. Host specificity was quantified as the number of distinct host species associated with each virus. Viruses detected in a single host species were classified as single-host viruses, whereas viruses detected in multiple host species were classified according to their observed host range. The distribution of host associations among proposed viral genera and plant families was visualized using standard statistical and graphical procedures. Where host-range distributions were compared between monocot-associated and dicot-associated groups, differences were evaluated using the Wilcoxon rank-sum test. Because transcriptomic datasets are unevenly sampled across host taxa, raw detection counts were interpreted together with the number of unique hosts, BioProjects and other available metadata to distinguish repeated sampling from independent host associations.

### 2.14. Large-scale retrospective search using Logan

To assess the broader diversity and distribution of lispi-like viruses beyond the newly assembled sequences, publicly available RNA-sequencing libraries were screened using the Logan search platform [41]. The 87 newly identified lispi-like virus sequences, together with MSuV and the previously reported Disporopsis pernyi-associated lispi-like virus, were used as query sequences. Searches were performed against the publicly indexed RNA-seq library collection using the standard Logan search procedure and parameters. Positive libraries were retained when they contained significant sequence matches to one or more lispi-like virus queries according to the platform’s default detection criteria. For each positive library, associated metadata were retrieved where available, including host species, botanical family, geographic location, sequencing project, and library identifier. Duplicate library records were retained as individual sequencing observations but were distinguished from independent BioProjects when assessing sampling redundancy. The final dataset was used to summarize the number of positive RNA-seq libraries, unique host species, host families, and geographic locations. Detection frequencies were interpreted as observations of viral sequence presence in public sequencing libraries and were not treated as direct estimates of biological prevalence in natural populations.

### 2.15. Virus prevalence and host-range metrics

To account for differences in transcriptomic sampling intensity among host systems, a normalized Virus Prevalence Score (VPS) was calculated for viruses detected through the Logan search as: [VPS = \frac{\mathrm{number\ of\ unique\ host\ species}}{\mathrm{number\ of\ positive\ RNA!-!seq\ libraries}} \times 100]. The VPS was used as a descriptive normalization of the relationship between host diversity and the number of positive libraries. Raw library counts, unique host counts and VPS values were considered jointly because each metric captures a different aspect of the available transcriptomic evidence. The number of independent BioProjects associated with each virus was additionally recorded to evaluate library redundancy and the extent to which repeated sequencing of the same biological system could inflate apparent detection frequency. Tissue-associated detections were classified according to the tissue metadata provided with the corresponding sequencing libraries. Tissue categories were standardized where possible, and the relative contribution of leaves, flowers, roots, stems, seeds, and whole-plant samples was summarized descriptively.

## 3. Results and Discussion

### 3.1 Unveiling plant-associated lispi-like viruses

Recent findings on the only two plant-associated lispi-like viruses identified thus far indicate that these viruses do not induce discernible symptoms in their natural hosts [28–29]. This phenomenon has also been reported for other negative-sense plant viruses, such as varicosaviruses [42], cytorhabdoviruses [11] and nucleorhabdovirus [43], suggesting that additional plant-associated lispi-like viruses might remain to be discovered in datasets derived from asymptomatic plants, where viral presence is not typically anticipated.

The advent of the powerful Serratus platform [8] has revolutionized the mining of this resource, enabling rapid and large-scale screening that would otherwise be prohibitively labor-intensive and time-consuming [11–12]. Leveraging this tool, we conducted the most extensive search to date on plant-associated lispi-like viruses RdRp sequences. This effort led to the identification and assembly of full coding regions of 87 novel lispi-like viruses, including several variants associated with different plant hosts (Table 1), increasing the number of known viruses in this group by 40X. Moreover, our findings significantly clarify their taxonomical position among the order *Mononegavirales*. These 87 viruses were associated with 75 plant hosts, where most of the apparent host plants are herbaceous dicots (57/75), while 18 hosts are monocots (Table 1). Thus, these viruses likely have a host adaptation trajectory leading to preferentially infecting herbaceous dicots during their evolution.

### 3.2 Genomic properties of lispi-like viruses

In contrast to other plant-associated mononegaviruses, such as the cytorhabdoviruses [11], among others, which display diverse genome organizations with many members having unique genome architectures [11, 44] all novel lispi-like viruses display four genes in a single genomic organization 3′-P1(N)-P2–P3-L-5′ (Table 1; Figure 1A); which was first reported for MSuV [28]. This genomic conservatism suggests a homogeneous evolutionary trajectory, and its linking to the family *Lispiviridae* tempts us to speculate that this novel group of viruses evolved from arthropod-infecting viruses, and lost the G gene during its plant adaptation. This hypothesis is supported by the seed-transmission nature of MSuV [28], suggesting that these viruses are transmitted vertically. Further studies should focus on their potential vector, if any, and its mode of transmission.

**Figure 1.**
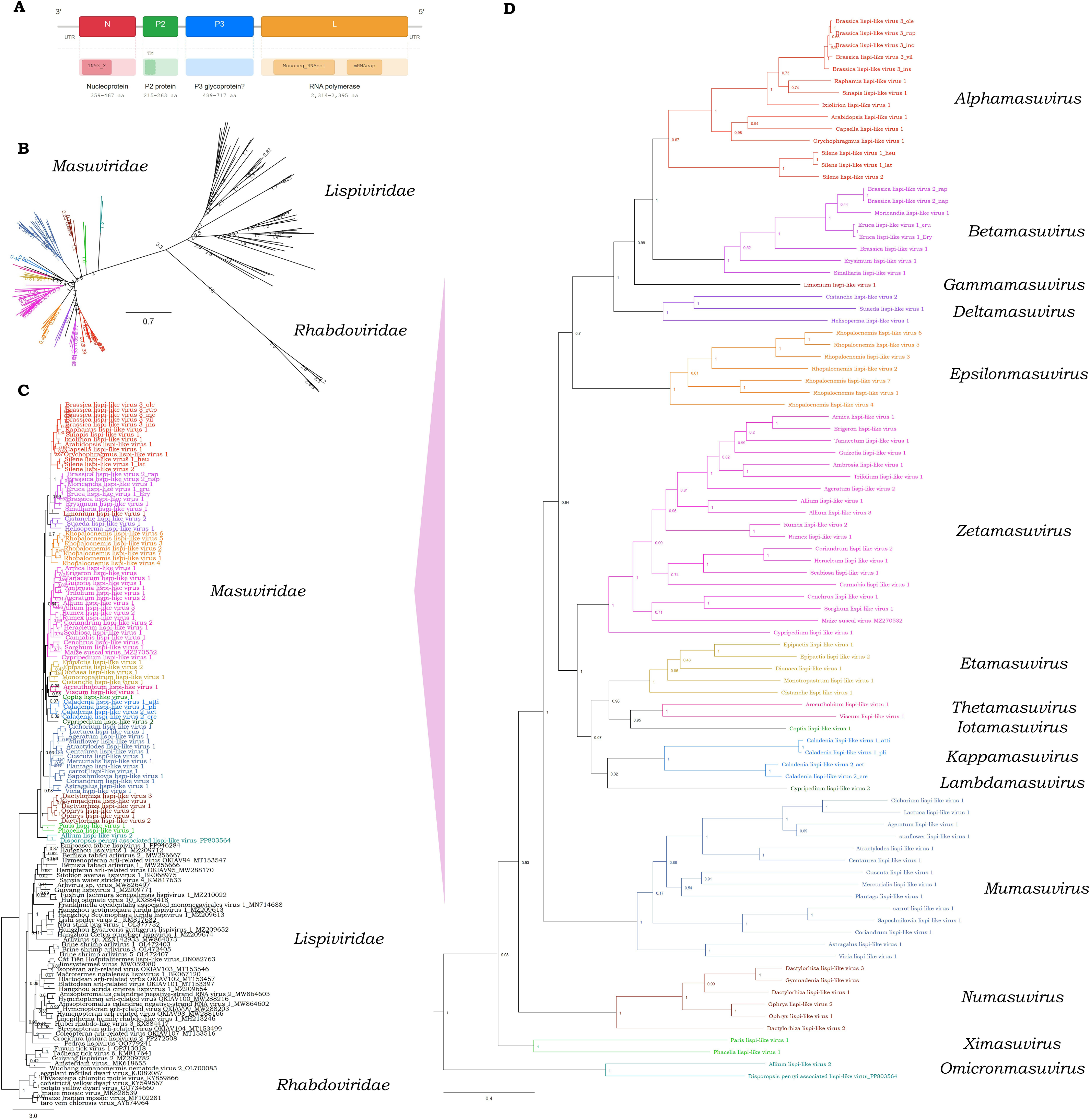
A. Genomic organization of the viruses identified in this study. B. Maximum-likelihood phylogenetic tree based on amino acid sequence alignments of the complete L gene of all reported so far and in this study constructed with the WAG + G + F model. The scale bar indicates the number of substitutions per site. Bootstrap values following 1,000 replicates are given at the nodes, but only the values above 50% are shown. Rhabdoviruses were used as outgroup.

The first ORF encodes the putative nucleocapsid, which is the protein commonly encoded by the first ORF in mononegaviruses [45]. The size of N proteins throughout the novel lispiviruses ranged from 359 to 467 aa (Table 1), which is similar to the size range reported for counterparts from for plant-associated mononegaviruses [26]. Moreover, BLASTP searches showed MSuV N protein as the best hit for each encoded N protein, but sharing a low identity value (Table 1). The second ORF encodes a protein, named as protein 2 (P2), and its size ranged from 215 to 263 aa. No similarity hits were found for this protein, even with relaxed parameters. Moreover, no conserved functional domains were identified in this protein, and any distant hit was found using HHblits or FoldSeek. Thus, further studies should be focused on the functional characterization of this protein to gain fundamental insights about its putative function. The third ORF encodes a protein, named as protein 3 (P3), and its size ranged from 489 to 717 aa. As with P2, no similarity hits nor conserved functional domains were found for this protein. The fourth ORF encodes the putative polymerase protein (L), and its size ranged from 2314 to 2395 aa, which is within the size range reported for L proteins from plant-associated mononegaviruses [26]. Moreover, BLASTP searches showed MSuV and Disporopsis pernyi associated lispi-like virus L proteins as the best hit for each encoded L protein, but sharing a low identity value (Table 1); and lispivirus L proteins were the following best hits with identities below 30%. The Mononeg_RNA_pol; Mononeg_mRNAcap and paramyx_RNAcap conserved domains were identified in each encoded L protein. These domains are conserved among the L proteins encoded by every mononegavirus [46].

The gene junctions of MSuV and of all newly identified viruses share a conserved, *Mononegavirales*-type architecture (Table 2). At the 3′ end of every gene, the mRNA-sense sequence terminates in a uridine-rich tract of five to seven residues preceded by an AU-rich segment (most commonly 3′-AU[C/A/U]UUUUUU), the expected signal for polymerase stuttering and mRNA polyadenylation in non-segmented negative-strand RNA viruses [46]. This termination signal is separated from the start of the downstream gene by an invariant trinucleotide intergenic spacer, most frequently CAC and less often CGC, and the downstream mRNA begins with a conserved trinucleotide, predominantly UCU or UCC. The three elements, terminator, spacer and initiator, are conserved both among the masuviruses themselves and relative to MSuV, and their organisation parallels the compact, highly conserved gene junctions reported for other plant-associated mononegaviruses [11, 42]. Moreover, MSuV genome has complementary ends; which is a common feature of mononegavirus [46]. Hence, the presence of a canonical U-rich termination/polyadenylation signal, a short conserved intergenic spacer and a conserved mRNA-start motif across the family provides independent, transcription-level support for the placement of this group within the order *Mononegavirales*, complementing the conserved genome organisation (Figure 1A) and the L-protein phylogeny (Figure 1B–D). The inclusion of plant lispi-like viruses within the *Mononegavirales* is further supported by morphological properties observed for MSuV, which possesses rod-shaped virions [28], similar to those found for other plant-associated mononegaviruses [26].

### 3.3 Evolutionary insights on masuviruses

Pairwise aa sequence identities between L proteins belonging to the same species ranged from 88.08 % to 99.45%. On the other hand, the pairwise aa sequence identities of L proteins across distinct species showed a great variation ranging from 29.92% to 79.98%, suggesting that there may still be an unknown amount of “virus dark matter” within plant-associated lispi-like viruses.

The phylogenetic analysis based on the L protein aa sequence showed that the 78 novel viruses, along with MSuV and Disporopsis pernyi associated lispi-like virus were grouped into a distinctive major group where four major clades could be distinguished (Figure 1B), and this group of viruses was linked to those members belonging to the *Lispiviridae* family (Figure 1B). This phylogenetic relationship indicates a common evolutionary history for this newly described group of viruses, where four clades of viruses showed distinct evolutionary trajectories.

Based on the phylogenetic insights and the observed genetic distance of the newly identified viruses we tentatively propose an 80% aa sequence identity of the L protein as the threshold for species demarcation in each one on newly proposed genera, the 78 members, for which the complete coding-sequence is available.

The phylogenetic analysis reveals that these plant-associated lispi-like viruses form a distinct monophyletic clade that diverges from the established family Lispiviridae, which comprises exclusively invertebrate-infecting viruses. This fundamental ecological transition between invertebrate and plant hosts, coupled with the substantial phylogenetic distance observed in RdRp-based trees, supports the establishment of a new viral family. The evolutionary trajectory evident in our analyses suggests an ancient host-switching event probably from arthropods to plants, followed by extensive diversification within plant hosts. This pattern of host realm transition accompanied by phylogenetic divergence parallels established criteria for family-level taxonomic distinctions in RNA virus classification. We therefore propose the family name *Masuviridae* (derived from **Ma**ize **su**scal **vi**rus, the first reported member of this lineage) to accommodate these plant-infecting viruses. The designation reflects both the historical precedence of maize suscal virus as the founding member and follows established viral nomenclature conventions. This taxonomic framework recognizes the fundamental biological differences between the invertebrate-associated *Lispiviridae* and their plant-associated sister group, while acknowledging their shared evolutionary origin within the broader lispi-like virus supergroup.

### 3.4 Structural Analysis of P1 Nucleocapsid Protein

Computational structural modeling of P1 nucleocapsid proteins from 87 lispi-like virus sequences revealed a canonical two-domain architecture characteristic of negative-strand RNA viruses [47], comprising an N-terminal domain (NTD), flexible interdomain linker, and C-terminal domain (CTD) (Figure 2). This bipartite organization mirrors the conserved "5H+3H" structural motif observed across the Negarnaviricota phylum [48–49], where nucleocapsid protein subunits are parallelly aligned along the RNA genome that is sandwiched between two domains composed of conserved helix motifs. AlphaFold2-based predictions showed high confidence for core functional domains (pLDDT >90) with excellent structural superposition against six experimentally determined nucleoprotein structures (PDB IDs: 6JC3, 4XJN, 1N93, 5WKN, 7EXA, 7NT5), while lower confidence in terminal regions (pLDDT 50-70) reflected expected intrinsic disorder in regulatory domains.

**Figure 2.**
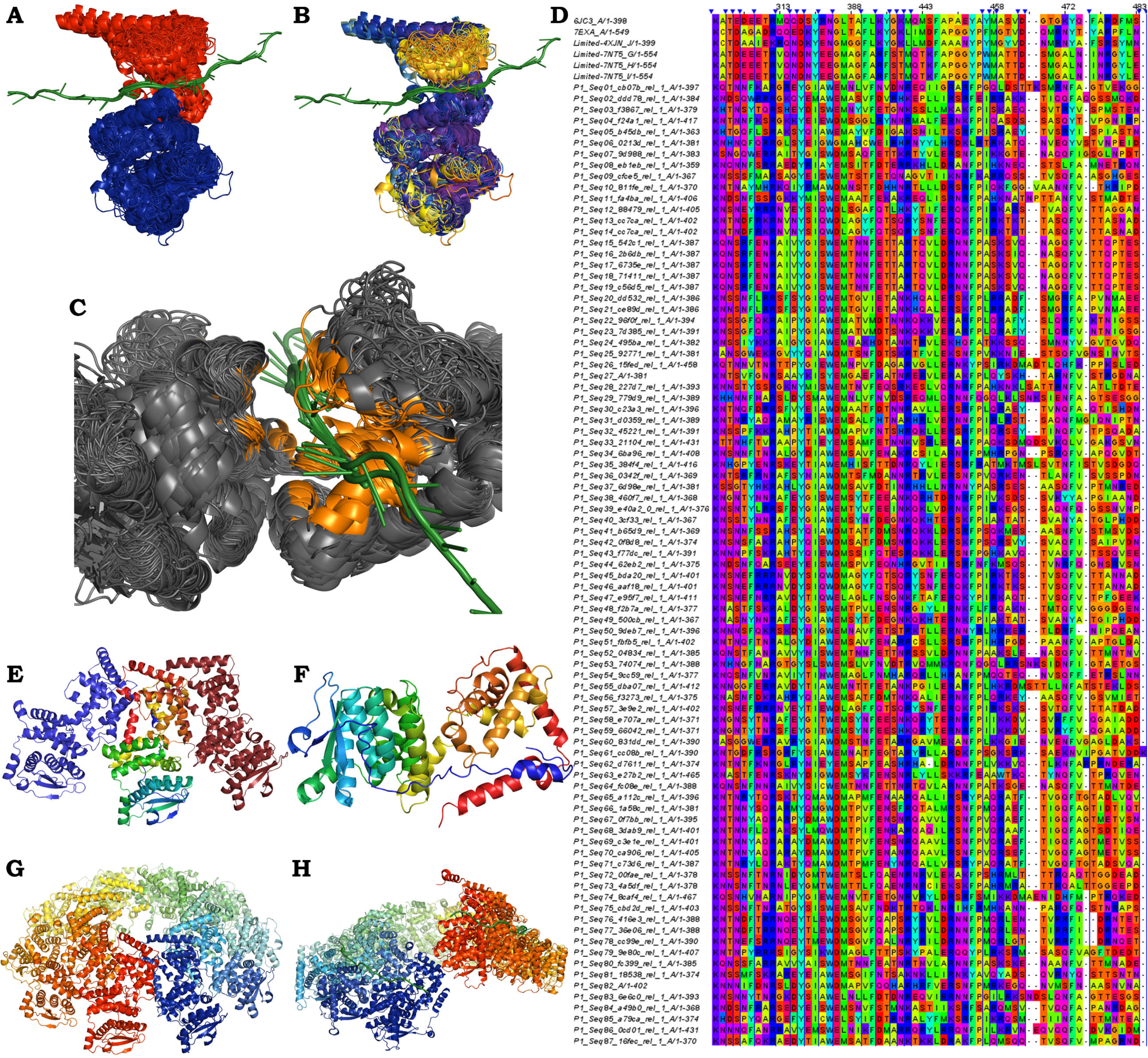
Structure of the P1 Nucleocapsid across Masuviridae shows consistent 5H+3H structure containing groove with 3 up and 3 down nucleotides protected by a lid. (A-C) Structures of P1 superimposed by (A) domain [blue Nterm, red Cterm], (B) pLDDT from purple [100%] to red [0%], and (C) RNA binding residues in orange and shown as an (D) MSA. (E) Structural trimer of P1 shown with monomers on the sides in blue and red while the central P1 shown in spectrum from Nterm [blue] to Cterm [red] which is also shown in (F) profile. (G) 10mer of P1 forming a circle and (H) 13mer CryoEM structure of 7NT5 [7] from Nipah virus.

The NTD exhibits a complex α/β architecture (β1-L1-α1-L2-α2-L3-α3-L4-β2-L5-α4-L6-α5-L7-α6-L8-α7-L9-α8-L10-α9-L11-α10) with critical RNA-binding residues showing exceptional conservation: K164 in loop L7 (100% conservation), R178 in helix α6 (99% as Arg/Lys), and Y242 in helix α10 (94% conservation). The interdomain linker contains highly conserved residues essential for RNA binding, including W249 (89% conservation) for aromatic stacking interactions and E251 (58% Glu, 34% Asp) for electrostatic RNA backbone contacts. The all-α-helical CTD (α1-L1-α2-L2-α3-L3-α4-L4-α5-L5-α6-L6-α7-L7-α8-L8-α9) features universally positively charged residues R299 (67% Arg, 26% Lys) and R312 (82% Arg, 12% Lys) for RNA phosphate interactions, alongside aromatic residues F257 (84%) and F315 (84% Phe, 15% Tyr) for nucleobase recognition.

This conservation pattern parallels observations in related virus families where no consensus interactions with the bases of the encapsidated RNA were identified among all reported nucleocapsid-like particle structures [47–48], suggesting that nucleocapsid proteins must accommodate all possible RNA sequences. The amino acid substitution patterns follow chemically conservative changes (Phe↔Tyr, Arg↔Lys), indicating that functional constraints, rather than phylogenetic relationships, drive sequence evolution at RNA-binding sites. Notably, regions α8, L7, and L8 in the CTD show high structural variability, suggesting adaptive mechanisms for accommodating diverse RNA sequences across different lispi-like viruses, similar to the protein plasticity enabling non-sequence specific interactions observed in Nipah virus nucleocapsid assembly [50] (Figure 2).

Structural analysis reveal that N protomers encapsidate RNA following the "3-bases-in, 3-bases-out" geometric arrangement with six nucleotides per protein monomer, consistent with paramyxovirus nucleocapsid assembly [50]. This packaging model predicts 1,500-2,000 N protomers are required for complete genome encapsidation (9,000-12,000 nucleotides), representing highly efficient packaging that maximizes RNA protection while minimizing protein investment. The proposed helical ribonucleoprotein complex forms through inter-protomer contacts creating continuous RNA-binding channels, enabling accommodation of variable genome lengths while maintaining consistent structural interactions (Figure 2).

The universal conservation of critical residues (K164, R178, W249, R312) demonstrates strong selective pressure to maintain specific RNA-protein contacts essential for genome packaging, aligning with the hallmark characteristic of negative-strand RNA viruses where genomes never exist as free RNA but are always assembled with nucleoproteins to form highly stable nucleocapsids. The near-universal conservation of W249 for RNA stacking and the conserved positive charge distribution (R178, R299, R312) present attractive targets for broad-spectrum antiviral development, as nucleocapsid-disrupting agents targeting these interactions could demonstrate pan-lispi activity.

While computational predictions show excellent agreement with experimental structures for core domains, experimental validation is still required to confirm those predictions. The low-confidence terminal regions likely reflect genuine intrinsic disorder consistent with regulatory functions, and static models cannot capture dynamic conformational changes during RNA binding and assembly. The proposed "3-bases-in, 3-bases-out" packaging model, while consistent with related nucleoproteins, represents one interpretation requiring validation through cryo-electron microscopy of complete nucleocapsid particles. Alternative packaging arrangements, including asymmetric RNA binding or variable stoichiometry models, should be considered alongside experimental mutagenesis targeting highly conserved residues to definitively establish structure-function relationships and validate the helical assembly mechanism proposed from our structural analysis.

### 3.5 Structural Analysis of P2

#### Structure-based functional annotation

To probe the function of the P2 protein, whose primary sequence is too divergent for reliable annotation by conventional sequence-homology search, we generated AlphaFold2 models for all 87 lispi-like virus P2 sequences and queried each predicted structure against the Protein Data Bank and the AlphaFold Database using Foldseek. This structure-based strategy exploits the fact that protein folding is considerably more conserved than the aa sequence [51], allowing remote homologies to be detected where the sequence signal has decayed [33].

The dominant outcome was an absence of interpretable annotation. Sixty-one of the 87 models (70%) returned no usable functional assignment: 30 matched only proteins that are themselves annotated as “uncharacterized”, 17 produced no significant structural hit, and 14 yielded matches too inconsistent or too low in confidence to interpret (Figure 3). P2 therefore belongs largely to the poorly characterized “dark” fraction of the viral proteome, in sharp contrast to the strong, internally consistent annotation obtained for the putative nucleocapsid from the same isolates.

**Figure 3.**
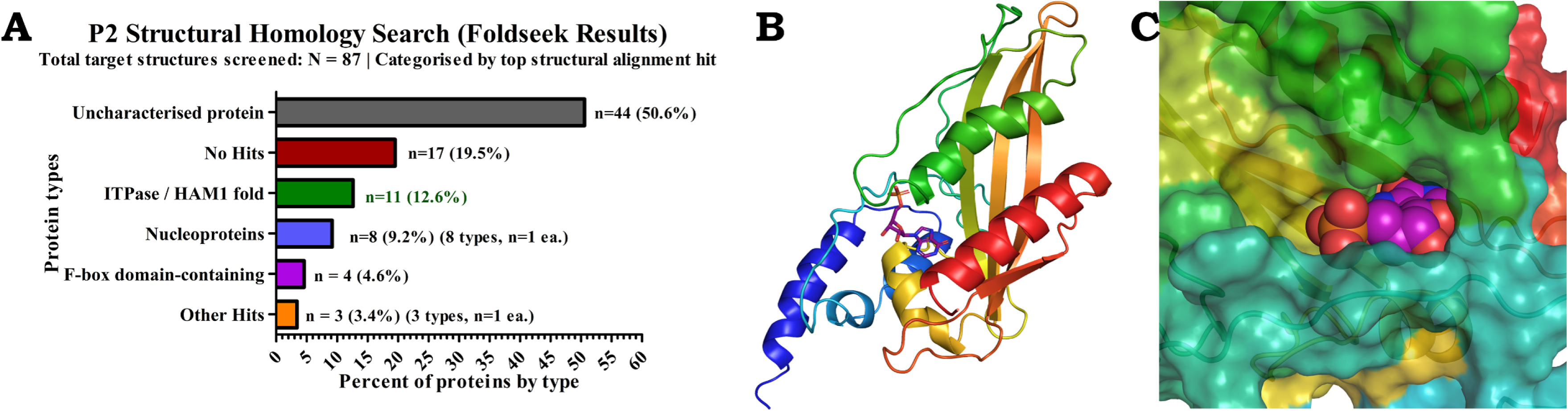
P2’s diversity (13.9% median identity) leads to only 11 structures labelled as ITPases. (A) P2 Foldseek results across 87 MuSV. Structure of P2 in ITPase protein structure shown with IMP [purple] in the catalytic pocket (B) as a cartoon and (C) with surface structure coloured in spectrum Nterm [red] to Cterm [blue].

Of the 26 models (30%) that did return a putative assignment, the single most frequent was inosine triphosphate pyrophosphatase (ITPase; Maf/HAM1 family), recovered for 11 sequences, roughly 42% of all annotated models and 13% of the full set. The remaining assignments were largely singletons spanning nucleotide- or nucleic-acid-associated activities (an ATP/GTP-binding protein, a cytosine/adenosine deaminase, GTPases, a valyl-tRNA ligase, a ribonuclease, a putative RNA-binding protein, and a zinc-finger), together with an F-box-related group and a small number of non-nucleotide annotations. In all, approximately three-quarters of the interpretable hits (19 of 26) corresponded to functions that act on nucleotides or nucleic acids; the dominant and most coherent signal is nucleotide metabolism, specifically purine-pool sanitization, rather than direct nucleic-acid binding.

The predicted P2 architectures were themselves structurally heterogeneous. Most models adopted a mixed α/β fold organized around a small central β-sheet (commonly two to five strands) flanked by α-helices; a substantial minority were entirely α-helical; a smaller subset displayed two distinct β-sheets; and four models (sequences 22, 33, 35 and 66) showed no discernible secondary-structure organization. The recurrent mixed α/β topology corresponds to the canonical architecture of the ITPase/HAM1/Maf superfamily, whereas the all-α subset is reminiscent of the distinct all-α (MazG-type) NTP-pyrophosphatase superfamily; both are among the four recognized structural classes of nucleotide “house-cleaning” enzymes [52]. For the ITPase-annotated models, qualitative inspection of cavity geometry, not automated docking, indicated a pocket dimensionally complementary to inosine that would sterically exclude the exocyclic N6 amino group of adenosine while leaving room for xanthine, a pattern mechanistically coherent with ITPase substrate selectivity (Figure 3). We stress that this is a static, qualitative observation presented as a structural prediction rather than evidence of catalytic activity.

#### Sequence-level properties and independent tests of the ITPase signal

The 87 P2 proteins form a single, full-length protein family: all begin with methionine, and their lengths occupy a narrow window (211–273 residues; median 231). Despite this coherence, the family is extraordinarily divergent, with a mean pairwise amino-acid identity of 13.9% (median 12.0%; range 3.1–99.6%), a value below the twilight zone of reliable sequence-based homology inference [53]. A multiple-sequence alignment recovered only a vestigial conserved core: of 217 well-populated columns, just 17 are conserved in at least half of the sequences and only four exceed 70% conservation, comprising fold-scaffold residues including a near-invariant tryptophan and lysine and a few buried hydrophobic and Gly/Pro positions. The heterogeneous structural annotations therefore reflect divergence and prediction noise within one protein family rather than a mixture of unrelated proteins.

Against this backdrop we tested the ITPase assignment with the most sensitive sequence-level methods available. Profile hidden Markov models for the Pfam Ham1 (PF01725) and Maf (PF02545) families [54–55] detected the three reference ITPases used as positive controls, human ITPA, *Escherichia coli* RdgB, and *Saccharomyces cerevisiae* Ham1, at high significance (176–223 bits; E ≤ 1 × 10⁻⁵⁴), yet returned no hit (E < 10) for any of the 87 P2 proteins, including all 11 that Foldseek had flagged as ITPase. Consistent with this, the flagged proteins shared only 4–10% identity with the reference ITPases in direct alignment, and the catalytic lysine that contacts the substrate triphosphate (human ITPA K19, equivalent to UCBSV Ham1 K38; 56) was conserved in just 2 of the 11. The ITPase signal is thus confined to the structural level and is not corroborated by sequence or profile homology.

The flagged proteins also do not constitute a coherent sequence lineage: their mean within-group identity (18.6%) is only marginally above their identity to the rest of the family (14.1%), and they disperse across a sequence-similarity clustering rather than forming a clade. They are, however, structurally self-consistent, with 10 of the 11 falling in the mixed α/β class. Physicochemically the family is near-neutral (mean isoelectric point ≈ 7) and slightly hydrophobic, and is not enriched in basic residues; the flagged subset is indistinguishable from the remainder on these measures. This composition is consistent with a globular enzyme and argues against a primary, basic-residue-driven nucleic-acid-binding role. Finally, the four models lacking discernible secondary structure are compositionally ordinary globular proteins by the charge–hydropathy criterion [57] rather than predicted to be intrinsically disordered, indicating that these cases most plausibly reflect low-confidence prediction rather than genuine disorder (Figure 3).

Mapping each isolate to its host yielded a broad plant range dominated by Brassicaceae, Asteraceae, Orchidaceae and Apiaceae and including several parasitic or mycoheterotrophic plants. Structural class showed no association with host family, and the 11 ITPase-flagged viruses spanned seven plant families with no representative of the *Euphorbiaceae*, the family of all four RNA viruses previously reported to encode a Maf/HAM1 domain [56]. The single *Euphorbiaceae*-infecting isolate in our set (*Mercurialis* lispi-like virus 1) returned no structural hit for P2. The host-linked pattern that characterizes the cassava system is therefore not reproduced here. We further note several tight within-species strain clusters (in the *Brassica*, *Caladenia*, *Eruca* and *Silene* virus groups); these introduce statistical non-independence and were treated as a caveat for any host-correlation or selection analysis.

The most prominent and internally consistent structural signal to emerge from this analysis is the recurrent assignment of a subset of P2 proteins to the ITPase (Maf/HAM1) family. ITPases are house-cleaning enzymes conserved across all domains of life that hydrolyze deaminated, mutagenic purine nucleotides, chiefly ITP, dITP, XTP and dXTP, to their monophosphates, preventing their misincorporation into RNA and DNA and thereby limiting mutation [58]. Their presence in RNA-virus genomes is rare: plant RNA viruses are under strong selection for compact genomes, and to date ITPase/Maf-HAM1 domains have been reported in only four RNA viruses, all infecting *Euphorbiaceae*, three potyvirids (cassava brown streak virus, Ugandan cassava brown streak virus, and euphorbia ringspot virus) and one secovirid, cassava torrado-like virus [56]. A genuine ITPase-like P2 in lispi-like viruses, if confirmed, would represent a phylogenetically independent and far broader occurrence of this unusual viral accessory function, which is precisely why the evidential basis for the assignment must be weighed carefully.

Experimental work on the cassava brown streak viruses provides the closest functional template. Their Ham1 proteins possess genuine *in vitro* ITPase activity with a preference for the non-canonical substrates dITP and XTP, act as determinants of necrotic symptoms [59], and are required for infectivity in cassava but dispensable in the model host *Nicotiana benthamiana*, a host-specific dependence attributed to the unusually high cytoplasmic concentration of non-canonical nucleotides in cassava, with the viral enzyme acting in partnership with the viral RNA-dependent RNA polymerase [56]. Under this model, a virus-encoded ITPase lowers the effective mutational load on the replicating genome, with the benefit, and therefore retention of the domain, contingent on the host’s nucleotide chemistry.

Our data, however, place clear limits on how far this template can be extended to P2. The structural evidence and the sequence evidence point in different directions. On the structural side, the ITPase assignment recurs across 11 models, those models reproduce the central-sheet-flanked-by-helices architecture of the ITPase/HAM1/Maf superfamily, and the predicted cavity is geometrically complementary to inosine, three observations that are mutually consistent. On the sequence side, the same proteins carry no detectable Ham1 or Maf profile signature, retain the catalytic lysine in only a minority of cases, do not form a coherent sequence group, and show no host association with *Euphorbiaceae*. The divergence of the family (mean identity ≈ 14%, below the twilight zone) means that the absence of a sequence signal is not by itself decisive, because fold is conserved more deeply than sequence [51]; a true but ancient homology could in principle have erased the profile signal. Even so, the negative result is informative, because the Ham1 profile readily recovers ITPases from bacteria, fungi and humans, and the failure to detect even a weak match across 87 proteins is a real constraint. The defensible conclusion is that nucleotide-pool sanitization remains the leading functional hypothesis for the ITPase-like subset, but that it now rests on structural evidence alone and should be presented as such.

Several alternatives merit explicit consideration. First, because Foldseek can return confident geometric matches between analogous but non-homologous folds, and because the flagged proteins are dispersed in sequence rather than forming a lineage, a fraction of the ITPase assignments may reflect structural convergence or model noise rather than homology. Second, the breadth of minor annotations and the structurally distinct all-α subset suggest that P2 may be a rapidly evolving, modular or multifunctional accessory locus rather than a single conserved activity. Third, even where an ITPase-like fold is genuine, catalytic competence is not guaranteed: the patchy conservation of the catalytic lysine is consistent with a structurally intact but catalytically degraded “pseudo-ITPase” in at least some isolates. These possibilities are not mutually exclusive, and distinguishing them is the central question for future work on P2. These conclusions rest on computational structure prediction and on sequence and profile search, and several limitations bound their interpretation. AlphaFold2 confidence was not uniform across models, and the no-fold and highly variable cases likely include low-pLDDT artefacts that inflate the apparent structural diversity; per-residue confidence should be reported alongside each model, and the four “no-structure” cases in particular reinterpreted as low-confidence predictions rather than disordered proteins. The inosine-pocket assessment is static and qualitative. The decisive remaining tests are therefore structural and biochemical rather than sequence-based: reading the residue structurally equivalent to ITPA K19 / Ham1 K38 directly from each predicted model; a profile–profile search (for example HHpred) as a final remote-homology check; and, definitively, an enzymatic assay of recombinant P2 against ITP and XTP versus canonical NTPs, ideally with a co-crystal or cryo-EM structure of an enzyme–substrate complex. A test of selection (dN/dS) on a codon alignment, and a reconciliation of the P2 phylogeny with the host and RNA-dependent RNA polymerase trees, would establish whether the ITPase-like variants are monophyletic and whether their retention tracks host taxonomy as in the cassava system; both require nucleotide and reference-tree data not analyzed here, and any such analysis should first collapse the within-species strain clusters identified above. Until then, the most defensible statement is that a structurally coherent subset of lispi-like virus P2 proteins resembles the ITPase/HAM1 family and presents an inosine-complementary cavity, making nucleotide-pool sanitization a plausible but unconfirmed function, while the majority of P2 sequences remain functionally uncharacterized and the family as a whole is too divergent for sequence-based annotation.

### 3.6 Structural Analysis of P3

The third major open reading frame (ORF3, encoding the protein termed P3) occupies the genomic position that, in the invertebrate-infecting *Lispiviridae*, encodes the viral glycoprotein, and a genome-wide alignment confirmed that all 87 viruses share this architecture. Consistent with extreme divergence, P3 returned no hits in BLASTp or PSI-BLAST searches, so it was characterised by structure prediction and by model-independent sequence analysis. AlphaFold2 modelling produced an essentially unstructured monomer, but assembly as a homotrimer yielded, for 54 of the 87 sequences, an elongated particle reminiscent of a class-I viral fusion protein in its post-fusion conformation, a globular head subtended by a long helical stalk, visually comparable to the post-fusion foamy virus envelope glycoprotein [60]. The confidence of these models was strongly non-uniform: the stalk helices modelled at moderate confidence (pLDDT ≈ 50–80) whereas the globular head fell below 40, that is, was effectively undetermined (Figure 4).

**Figure 4.**
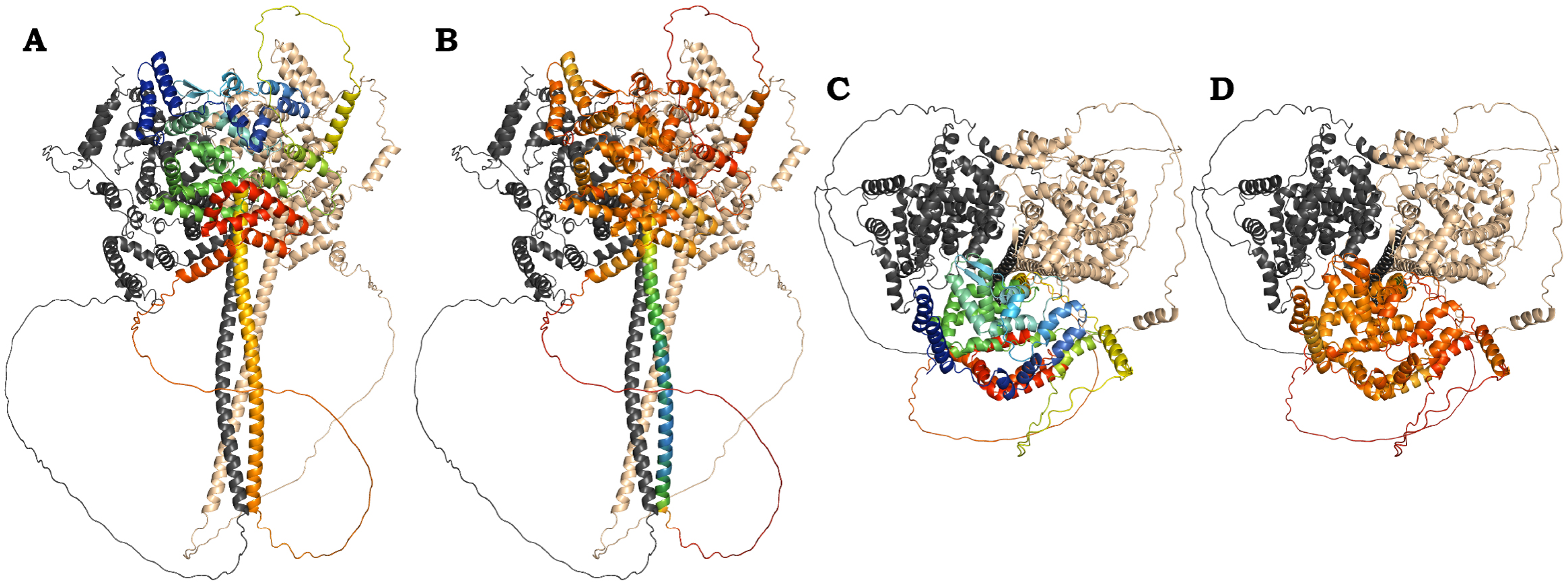
52/87 P3 proteins formed glycoprotein with a confident coiled-coil but an uncertain head region resembling post fusion proteins. (A) P3 trimer shown from (A-B) the side and (C-D) the top, with chain A of each denoting a spectrum of (A,C) Nterm [blue] to Cterm [red] or pLDDT confidence (B,D) from purple [100%] to red [0%].

At the sequence level, the 87 P3 proteins form a single, full-length family, all begin with methionine, and lengths occupy a coherent window (489–717 residues, mean 624), roughly 300 residues shorter than the foamy virus glycoprotein precursor. The family is nonetheless hyper-divergent, with a mean pairwise amino-acid identity of 13.3%, below the twilight zone of reliable sequence-homology inference [53], which accounts for the failed similarity searches. Against this low background one feature is conserved family-wide: every sequence carries a sustained heptad-repeat coiled-coil, and the coiled-coil register is positionally conserved across the alignment, with all 87 sequences in register at the most conserved column. This conserved coiled-coil is the element that forms the trimeric stalk in the models and is, by a wide margin, the most strongly constrained part of the protein (Figure 4).

The membrane-protein hallmarks expected of a surface glycoprotein are, however, absent at the family level. Combined signal-peptide and transmembrane-topology prediction (Phobius; 32) assigned a signal peptide to only 6 of 87 sequences, five of which constitute a single near-identical *Brassica* strain cluster, and a transmembrane segment to just one, so the large majority are predicted to be cytoplasmic, with neither an N-terminal secretion signal nor a C-terminal membrane anchor. The proteins are cysteine-poor (mean ≈ 5 residues, ≈ 0.7%), unlike the disulphide-stabilised ectodomains of characterised fusion glycoproteins, and N-glycosylation sequons occur at approximately the frequency expected by chance (≈ 3.7 per protein) with no sequon conserved across the family. A substantial fraction of each protein is predicted to be intrinsically disordered (mean ≈ 30% of residues, up to ≈ 68% in individual sequences), concentrated in the head and in long inter-domain loops, the same regions that modelled at low pLDDT.

Because P3 is encoded as a standalone cistron rather than within a polyprotein, any proteolytic maturation would have to be carried out in trans by host proteases. Scanning for plant-relevant cleavage motifs, the Site-1-protease/SKI-1 ortholog (SBT6.1/AtS1P), vacuolar legumains/VPEs, and, for completeness, furin-type sites (noting that plants lack the metazoan furin convertases), recovered no motif positionally conserved across the family, and no candidate located at the head–stalk junction ahead of a hydrophobic, fusion-peptide-like segment. When conservation was instead assessed within the 11 multi-member, host-coherent groups recoverable from this sparsely sampled, roughly 13–15-genus family (mean within-group identity 67%), cluster-level conserved sequons and S1P-type motifs did emerge, candidates that a family-wide scan necessarily discards. Their interpretation is bounded on two counts: the short S1P-type motif is frequently shared within a group by descent rather than by selection, and the relevant plant proteases, like N-glycosylation itself, act only on proteins that enter the secretory pathway, which here is restricted to the two signal-peptide-bearing lineages (*Brassica* and *Limonium*).

Selection analysis on a codon alignment confirmed that P3 is a genuine, functionally constrained gene. Family-wide, synonymous sites are saturated (proportion of synonymous differences ≈ 0.75), so global rate ratios are uninterpretable; estimated within low-divergence pairs, the protein is under clear purifying selection (dN/dS ≈ 0.10), and the constraint is spatially structured, the coiled-coil core being markedly more constrained than the head and loops (dN/dS ≈ 0.08 versus ≈ 0.24). Site-wise analysis using within-cluster comparisons classified 61% of testable codons as significantly purifying and none as diversifying. The variable head is therefore under relaxed purifying selection rather than positive selection, and shows no signature of the antigenic or receptor-driven diversification often seen in the receptor-binding heads of fusion glycoproteins.

Three lines of evidence place P3 as the plausible, if highly divergent, positional homolog of the *Lispiviridae* glycoprotein. It occupies the syntenic glycoprotein cistron across all 87 genomes; it assembles into an elongated homotrimer whose architecture resembles a post-fusion class-I fusion protein [60–61]; and its single conserved, family-wide structural element is a coiled-coil that forms the trimeric stalk, the heptad-repeat coiled-coil being the defining core of class-I fusion machinery [62]. The selection landscape reinforces this reading at the level of architecture, in that the coiled-coil core is the most strongly purifying-selected region while the head is variable, paralleling the conserved fusion core and variable head of characterised fusion proteins.

Against this, the features that define a *matured surface* glycoprotein are absent family-wide. Class-I fusion glycoproteins are synthesised with an N-terminal signal peptide, anchored by a C-terminal transmembrane domain, N-glycosylated on a disulphide-stabilised ectodomain, and proteolytically primed by a host protease [61–62]. P3, as predicted from sequence, has none of these as a conserved property: no signal peptide or transmembrane anchor in the large majority, few cysteines, chance-level glycosylation, and no family-wide cleavage site. Because the family is so divergent that synonymous sites are saturated and identity lies below the twilight zone, the absence of these signals cannot by itself exclude a fusion glycoprotein, divergence can erode or relocate signal peptides, anchors and cleavage sites, but neither do the data positively support a canonical, membrane-displayed, glycosylated, primed glycoprotein. The defensible reading is that the conserved coiled-coil and the trimeric model are real and fusion-protein-like, whereas the maturation and membrane-targeting apparatus expected to accompany them is not detectable at the family level.

The strength of the structural impression also warrants caution about its origin. AlphaFold-Multimer readily generates plausible, symmetric, elongated coiled-coil assemblies, and a model combining a confident central coiled-coil with a low-confidence peripheral domain is a recognised failure mode rather than necessarily a biological structure [38]. Here the head modelled below 40 pLDDT, effectively unmodelled, and three independent measures (pLDDT, predicted disorder, and relaxed selection) agree that this region is poorly determined. The post-fusion resemblance therefore rests on the conserved coiled-coil, the synteny, and the trimerisation, none of which establishes a fusion function on its own. Quantitative structural comparison, for example Foldseek or TM-align against experimentally determined post-fusion fusion proteins [33], together with interface-confidence metrics (ipTM, predicted aligned error) would be required to test the resemblance properly.

The diversity of the family shapes how its functional features should be read. With members spanning on the order of 13–15 prospective genera, candidate functional sites are not expected to be conserved across all 87 sequences, and indeed glycosylation sequons and S1P-type cleavage motifs are conserved within genera even though none is conserved family-wide, a pattern that a whole-family conservation test would erroneously read as absence. Whether any of these represents genuine, used machinery is gated by subcellular localisation: only the two secretory lineages could be glycosylated or processed by secretory plant proteases, so genus-level processing, if it occurs, is likely lineage-specific rather than a family-wide property. Beyond a divergent fusion protein, the data remain compatible with alternative roles for a conserved trimeric coiled-coil protein at this genomic position, such as a structural or scaffolding function; and the absence of diversifying selection argues specifically against a classical antigenic receptor-binding head, though not against a structured head that simply was not modelled.

Several limitations bound these conclusions. The structural inferences rest on AlphaFold models whose interface confidence was not evaluated here and whose head is undetermined; the topology, glycosylation and cleavage predictions are sequence models that may miss non-canonical determinants in a divergent viral protein; selection was estimated by Nei–Gojobori counting [40] rather than by a maximum-likelihood site model, so the count of purifying sites is a lower bound and the absence of diversifying sites should be confirmed per genus with a likelihood method [63]; and cluster-level conservation partly reflects shared ancestry. Resolving the identity of P3 will require experiment, N-terminal sequencing or mass spectrometry of virion-associated P3 to detect maturation, glycosidase and membrane-fractionation assays to test glycosylation and membrane association, and ultimately a determined trimer structure. Until then, the data read most defensibly as a conserved, trimer-forming, coiled-coil protein occupying the glycoprotein cistron and under strong purifying selection on its core, carrying a structurally undetermined, disordered, lineage-variable head, and lacking the family-wide hallmarks of a matured surface glycoprotein.

### 3.7 Structural Analysis of P4

#### The largest lispi-like virus ORF encodes an L-protein-like RdRp

The fourth open reading frame encodes the large (L) protein, hereafter also P4, which is by a wide margin the largest cistron in the lispi-like virus genomes; its 87 translated products are strikingly uniform in length (2,314–2,402 aa; mean 2,348 aa, s.d. 18 aa). This size and the collinear arrangement of conserved blocks are characteristic of the multidomain L protein of the *Mononegavirales*, which integrates the RNA-dependent RNA polymerase (RdRp), the polyribonucleotidyltransferase (PRNTase) capping activity, and a cap methyltransferase within a single polypeptide [39, 46, 64]. Because the RdRp is the single most conserved and most taxonomically diagnostic protein across RNA viruses, and the primary character used by the ICTV to delimit *Mononegavirales* families [25, 65], we examined P4 both structurally and at the sequence level to test its identity and its catalytic competence.

#### The structural model recovers a canonical RdRp core but not the accessory domains

An AlphaFold2 model of a representative P4 [37] was compared with the cryo-EM structure of the vesicular stomatitis virus (VSV) L protein (PDB 5A22; 39), the prototypical *Mononegavirales* polymerase, using structural superposition and Foldseek [33] (Figure 5D–F). The comparison resolves cleanly into two regimes. The N-terminal RdRp module of P4 superimposes on the VSV RdRp core (Figure 5E–F, core in red) and reproduces the canonical right-hand polymerase architecture, with clearly delineated fingers, palm and thumb subdomains (Figure 5G). Within the palm, the three catalytic palmprint motifs A, B and C [64, 66] are present and correctly positioned (Figure 5H), and the key metal-coordinating and template-contacting residues project into the active-site cleft in the expected geometry (Figure 5I). By contrast, the domains that flank the RdRp in a complete L protein, the PRNTase/capping domain, the connector domain, the methyltransferase and the C-terminal domain (Figure 5D), did not superimpose confidently onto their VSV counterparts and were not resolved as discrete, VSV-like folds in the model. The structural evidence therefore supports a bona fide RdRp core embedded in a large L-like protein whose accessory modules could not be confidently modelled by homology.

**Figure 5.**
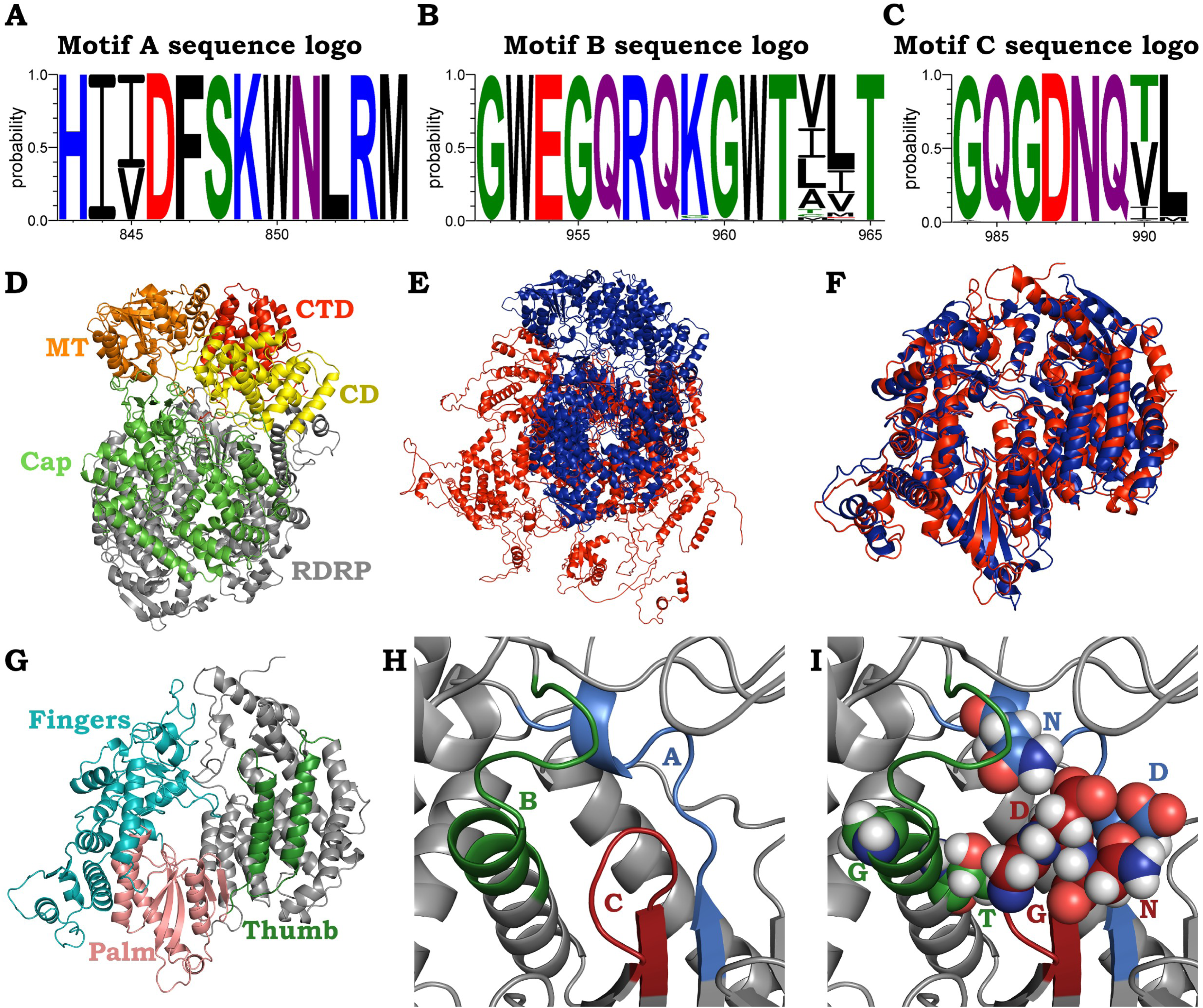
P4 of Masuviridae maintains core RDRP and catalytic palmprint but surrounding domains were not resolved. (A-C) Sequence logos of the 3 core motifs of RDRP. (D) Structure of VSV RDRP with all domains from 5A22 [8]. (E-F) Structure of P4 and the core RDRP [red] compared to VSV [blue]. (G) RDRP of P4 shown highlighting the Thumb, Palm, and Fingers of the core structure. (H) Zoomed view of the Palmprint motifs of P4 RDRP with (I) key residues highlighted.

#### P4 is markedly more conserved than the other lispi-like virus proteins, and its divergence is structured by host family

Across the 87-sequence alignment, P4 shows a mean pairwise amino-acid identity of 39.9% (median 38.5%, range 29.0–99.5%; Supp. Fig. S1A) [HARMONISE with §3.2, which reports the same L protein as 29.92–79.98% between species and 88.08–99.45% within species, adopt one identity computation and cross-reference]. This places P4 at a far higher level of conservation than the P3 glycoprotein-like product of the same viruses (≈13% mean identity; §3.5), as expected given the exceptional selective constraint on RdRp catalytic function [67]. The divergence is nonetheless substantial and consistent with family-rather than genus-level breadth. Pairwise identities are structured by host botanical family: sequences from viruses sharing a host family are appreciably more similar to one another (mean 50.5%) than to sequences from other host families (mean 38.8%), and the identity matrix resolves coherent within-family blocks, most conspicuously for Orchidaceae-, Brassicaceae- and Balanophoraceae-associated viruses (Supp. Fig. S1B).

#### Conservation is concentrated in the RdRp core, mirroring the structural result

The per-site conservation profile provides a direct sequence-level counterpart to the structural observation (Supp. Fig. S1C). Conservation is not uniform along P4: the palmprint-bearing N-terminal moiety, corresponding to the structurally resolved RdRp core, is significantly more conserved than the C-terminal moiety that harbours the capping, methyltransferase and C-terminal domains (mean normalised conservation 0.69 versus 0.49; Mann–Whitney U test, one-sided, p ≈ 2 × 10⁻⁶⁵). The three catalytic motifs sit within near-invariant columns (Shannon entropy ≈ 0.00–0.03 bits) against a genome-wide median column entropy of 1.92 bits, and 782 alignment columns are effectively invariant (entropy < 0.5 bits). Thus, the same partition seen structurally, a well-ordered, superimposable RdRp core versus poorly resolved accessory domains, is recovered as a quantitative divergence gradient in the primary sequence, with the accessory region evolving substantially faster than the core.

#### The catalytic palmprint is invariant and carries the Mononegavirales GDN signature

At single-residue resolution, the catalytic apparatus is completely conserved across all 87 sequences (Supp. Fig. S1D–F). The metal-coordinating aspartate of motif A is present in 87/87 sequences within an invariant DFSKW block, and the catalytic triad of motif C is G-D-N in 87/87 sequences, embedded in the pentapeptide QGDNQ that is invariant in 86/87 sequences (the single exception, an A-for-G substitution at the first flanking position, leaves the catalytic GDN intact). Motif B (block GWEGQRQKGW) is conserved in 83/87 sequences (95.4%), with the four variants being conservative substitutions that preserve the block framework. No sequence carries a GDD or SDD variant at motif C.

The identity of the motif-C triad is phylogenetically informative. Among viral polymerases the motif-C signature is group-specific: GDD in the RdRps of most positive-strand RNA viruses, SDD in the L proteins of segmented negative-strand RNA viruses, and GDN in the L proteins of the non-segmented negative-strand viruses of the *Mononegavirales* [68–69], where the GDNQ motif is functionally essential [70–71] and the QGDNQ pentapeptide is recognised as one of the longest invariant stretches of the polymerase domain [72]. The uniform GDN of P4 therefore places its polymerase squarely in the *Mononegavirales* catalytic class and is inconsistent with a segmented- or positive-strand origin. Together, the invariant motif-A aspartate and the invariant GDN of motif C reconstitute both catalytic aspartates of the two-metal-ion mechanism [64, 68], providing strong sequence support for a catalytically competent polymerase.

The convergent structural and sequence evidence establishes P4 as a functional-grade, *Mononegavirales*-type RdRp, while also delimiting the confidence of that assignment domain by domain.

#### The catalytic core is intact and its assignment is robust

The recovery of a canonical fingers–palm–thumb fold that superimposes on VSV L (Figure 5E– G), combined with the 100% conservation of the motif-A aspartate and the motif-C GDN across all 87 viruses (Supp. Fig. S1D–F), makes the identification of P4 as an active RdRp about as secure as sequence and modelling evidence allow. The concordance is not trivial: structure alone can be over-interpreted at low homology, and sequence motifs alone can be recovered in pseudogenised or defective polymerases, but their agreement here, a resolved active-site geometry occupied by an invariant, correctly spaced catalytic constellation, is difficult to reconcile with anything other than a bona fide polymerase. The GDN signature further fixes the catalytic class as *Mononegavirales* rather than segmented (SDD) or positive-strand (GDD); consistent with this, experimental substitution of GDN by GDD or SDD abolishes or severely impairs L activity in rabies virus and VSV [68], so the uniform GDN of P4 is the signature expected of a functional non-segmented negative-strand polymerase and supports the family’s proposed placement as a plant-associated lineage within the *Mononegavirales*.

#### The unresolved accessory domains most plausibly reflect divergence, not absence

The failure of the capping, connector, methyltransferase and C-terminal regions to superimpose on their VSV counterparts (Figure 5D–F) admits several explanations, and the sequence data help to discriminate among them. The first, that these domains are genuinely absent, is not supported: P4 is full-length and remarkably length-invariant (s.d. 18 aa over 87 sequences), leaving no room for large deletions of an entire capping or methyltransferase module. The second, that the region is intrinsically disordered, is also unlikely, because the C-terminal moiety retains appreciable sequence conservation (mean normalised conservation 0.49; Supp. Fig. S1C), well above what unconstrained sequence would show, indicating ongoing purifying selection rather than degeneration. The most parsimonious interpretation is the third: these are genuine but strongly divergent accessory domains whose remote homology to *Rhabdoviridae* fell below the confidence threshold of homology-based structural modelling, so that AlphaFold2 could neither template them on 5A22 nor place them by co-evolutionary signal. This interpretation is consistent with the pronounced core-versus-accessory conservation gradient (p ≈ 2 × 10⁻⁶⁵), itself the expected signature of a conserved catalytic core carrying faster-evolving peripheral modules. We therefore describe the accessory domains as unresolved rather than lost, and caution against inferring their absence from the model.

Several caveats constrain the present claims. First, functional competence is inferred, not demonstrated; the catalytic assignment remains a strong hypothesis pending experimental evidence, which for these uncultured, metagenomically defined viruses will require heterologous expression and in vitro polymerase or minigenome assays. Second, we did not test the accessory region for the diagnostic sequence signatures of its putative functions, for example the collinear PRNTase capping motifs A–E and the invariant HR of the capping active site [73–74], or the K-D-K-E tetrad of the 2′-O-methyltransferase [46]; confirming or refuting these would materially sharpen the interpretation of the unresolved domains and is a clear next step.

### 3.8 Genus-level classification and host associations

Based on phylogenetic analysis of RdRp sequences and pairwise identity thresholds, we propose the classification of *Masuviridae* into 15 genera designated *Alphamasuvirus* through *Omicronmasuvirus* following established nomenclature conventions (Figure 1.C-D). This taxonomic framework is supported by both molecular phylogeny and distinct ecological patterns in host plant associations.

The genus assignments were established using a combination of phylogenetic clustering and RdRp amino acid identity thresholds, with genera showing discrete monophyletic groups and identity values typically below 40% between genera. Within genera, virus species exhibited higher sequence conservation (>40% identity) while maintaining distinct host ecological niches. The largest genus, *Zetamasuvirus*, encompasses 19 virus species, while several genera (*Gammamasuvirus*, *Iotamasuvirus*, *Lambdamasuvirus*) are currently represented by single species, reflecting either restricted geographic sampling or genuine ecological specialization. Analysis of 89 confirmed virus-host associations across the 15 genera revealed striking patterns of host specificity that support the proposed taxonomic divisions. Most notably, several genera exhibit strong host plant family specialization. *Numasuvirus*, *Kappamasuvirus*, and *Lambdamasuvirus* are exclusively associated with *Orchidaceae*, with *Numasuvirus* showing the broadest orchid host range (6 viruses across 4 orchid species). Similarly, *Betamasuvirus* demonstrates complete fidelity to Brassicaceae (8 viruses), while *Alphamasuvirus* shows strong preference for this family (71% of associations). The most extreme case of host restriction occurs in *Epsilonmasuvirus*, where all seven virus isolates derive from a single host species, *Rhopalocnemis phalloides* (Balanophoraceae), suggesting either recent host colonization or highly specialized adaptation.

In contrast, *Zetamasuvirus* and *Mumasuvirus* exhibit broader host ranges spanning 9 and 6 plant families, respectively, with Asteraceae representing the most frequent host family overall (24 virus associations). This pattern suggests these genera may represent either ancestral generalist lineages or taxa with enhanced host-switching capabilities.

The observed host associations likely reflect multiple evolutionary and ecological factors including geographic sampling bias toward agricultural species, phylogenetic constraints on host compatibility, and vector specificity where applicable. The predominance of dicot hosts (78% of associations) and the concentration of viruses in economically important plant families (*Asteraceae*, *Brassicaceae*, *Orchidaceae*) may partly reflect detection bias in transcriptomic datasets. However, the genus-specific clustering of host associations indicates genuine biological constraints rather than purely technical artifacts.

These host association patterns provide additional support for the proposed genus-level classification and suggest that ecological specialization has been a significant driver in *Masuviridae* diversification. The combination of phylogenetic divergence and host specificity patterns observed here parallels those documented in other plant virus families where host adaptation and geographic isolation promote speciation and taxonomic diversification.

### 3.9 Viral diversity and prevalence patterns

We employed the Logan search tool (https://logan-search.org/dashboard) [41] to explore all publicly available libraries in order to search for the newly identified lispi-like virus sequences as well as MSuV and Disporopsis pernyi associated lispi-like virus. Our comprehensive data mining identified 1,536 public RNA-seq libraries containing lispi-like viral sequences across 134 host plant species, representing 11 plant families from 182 geographic locations worldwide. This substantial viral diversity, accumulated over nine years of sequencing data (2015-2024), demonstrates the power of retrospective bioinformatic mining to uncover previously unrecognized plant virus diversity hiding within transcriptomic datasets generated for other research purposes. Furthermore, it is tempting to speculate that substantial remaining diversity is still hidden; thus, a comprehensive sampling of global plant diversity could reveal hundreds of additional lispi-like viruses.

The application of a normalized Virus Prevalence Score (VPS), calculated as (Unique Hosts/Libraries) × 100, is key to understand the virus importance compared to raw detection frequencies (Supp. Fig. 2, Supp. Fig. 3). While Brassica lispi-like virus 2 dominated the raw library counts with 509 detections, VPS normalization revealed Astragalus lispi-like virus 1 as the most ecologically prevalent (VPS=163), infecting diverse hosts despite fewer absolute detections. This normalization accounts for sampling bias inherent in heavily studied agricultural systems and model organisms, providing a more accurate assessment of true host range breadth. This disparity between raw counts and normalized prevalence underscores that detection frequency often reflects research intensity rather than biological significance [75], a critical consideration for data-mining studies that must contend with highly uneven sampling across the plant kingdom.

The relationship between library abundance and host diversity revealed distinct ecological strategies among the identified viruses (Supp. Fig. 4). Brassica-lispi like viruses 2 and 3 exemplify specialists with narrow host ranges but high detection frequencies within their preferred hosts (18-20 organisms, primarily *Brassicaceae*). In contrast, viruses like the Astragalus lispi-like virus 1, despite moderate library counts, demonstrated broader host diversity with high detection robustness, suggesting true generalist capabilities rather than sampling artifacts. Raphanus lispi-like virus 1 occupied an intermediate position with both substantial library counts and moderate host diversity (14 organisms), potentially representing an emerging generalist lineage. This diversity of ecological strategies parallels patterns observed in other plant virus families, where host range evolution proceeds along multiple trajectories shaped by vector specificity and cellular compatibility, as well as geographic opportunity [76–77].

### 3.10 Host specificity and range constraints

Analysis of host specificity patterns revealed that strict host specialization predominates among lispi-like viruses, with 53 of 80 viruses (66%) restricted to single host species. This extreme specialization contrasts with the small number of generalists capable of infecting 4-12 different hosts (n=6 viruses, 7.5%). The negative correlation between sampling intensity and host specificity index suggests that apparent generalism may partially reflect detection bias, where those viruses sampled more intensively have greater opportunity to reveal additional hosts. However, the existence of true specialists even among heavily sampled viruses (e.g., Brassica lispi-like virus 3 with 172 libraries but restricted to Brassicaceae) indicates that biological constraints, not merely sampling limitations, define host range boundaries for the lispi-like viruses, such as has been reported for other plant viruses [77–78].

Moreover, the comparison of monocot-specific versus dicot-dominated viruses revealed no significant difference in host diversity where Dicot-dominated viruses (green, n=∼120) show broader distribution with median around 1-2 hosts and outliers extending to 20 hosts, whereas Monocot-specific viruses (coral, n=∼15) display narrower distribution centered around 1-2 hosts with maximum diversity of 5 hosts (Supp. Fig. 5). However, the Wilcoxon test (p=0.3636) (Supp. Fig. 5) indicates no statistically significant difference in host diversity between groups, suggesting that host range breadth is not determined by the monocot/dicot classification of primary hosts.

### 3.11 Library redundancy and sampling bias

The substantial discrepancy between total library counts and unique BioProjects for top-ranking viruses highlights a critical challenge in interpreting data-mining results (Supp. Fig. 6). Brassica lispi-like virus 2, with 509 libraries derived from only 72 independent studies, demonstrates how intensive sampling of specific experimental systems can inflate apparent prevalence. This redundancy particularly affects agricultural and model organism systems where researchers conduct time-series experiments, biological replicates, and multi-condition comparisons, all generating separate SRA entries that represent the same geographic population or experimental lineage [79].

Tissue tropism analysis revealed strong bias toward foliar sampling, with leaf tissue representing >75% of detections for most viruses (Supp. Fig. 6). This sampling artifact limits our ability to assess true tissue specificity, where viruses appearing leaf-specific may simply reflect that leaves are the predominantly sampled tissue [80]. The exceptions, such as Raphanus lispi-like virus 1 and Brassica lispi-like virus 3 with documented flower, root, stem, and whole plant detections, suggest these viruses either possess genuine systemic infection capability or have been subjects of more comprehensive tissue surveys. Seed-associated detections for Ageratum-lispi like viruses 1 and 2 and Arnica lispi-like virus 1, as well as flower-associated detections for Monotropastrum lispi like virus 1, Ophrys lispi like virus 1, Caladenia lispi like virus 1 and Gymnadenia lispi like virus 1(Supp. Fig. 6) raise questions about vertical transmission potential that cannot be addressed without targeted experimental follow-up. Other negative-sense plant viruses were reported to be seed-associated [81–83], and the lispi-like virus MSuV was found to be seed-transmitted [28], supporting the hypothesis that lispi-like viruses could be vertically transmitted.

### 3.12 Agricultural relevance and wild plant reservoirs

The agricultural relevance analysis revealed a dichotomy between crop-associated viruses and those circulating in wild plant populations. Brassica lispi-like viruses 1 and 2 showed >95% association with major crops despite limited host diversity, representing viruses of immediate agricultural concern that may cause unreported or subclinical impacts on crop productivity. In contrast, numerous lispi-like viruses detected exclusively in wild plants represent a largely unexplored reservoir of viral diversity with unknown agricultural risk. These wild plant viruses could potentially represent preadapted lineages capable of emerging into agricultural systems, similar to historical examples where crop domestication or agricultural intensification facilitated virus host shifts [75]. Raphanus lispi-like virus 1, with ∼75% crop association across 14 hosts, may exemplify an intermediate stage where viruses maintain circulation in both agricultural and wild populations, potentially using wild hosts as reservoirs for recolonization after crop rotations or seasonal cycles.

### 3.13 Geographic distribution

Global sampling coverage (Supp. Fig. 7) revealed strong geographic bias toward northern hemisphere temperate regions, particularly East Asia, Europe, and North America. China emerged as a hotspot with the highest virus richness (more than 15 viruses), likely reflecting both genuine biodiversity and substantial investment in plant genomics research. While sampling extended to 182 unique locations across 252 countries, density remained highest in regions with established agricultural research infrastructure and genomics capacity. This geographic bias has important implications for understanding global viral biogeography. Tropical and southern hemisphere regions remain undersampled despite harboring the majority of plant species diversity [84–85], suggesting substantial undiscovered viral diversity in these areas.

### 3.14. Implications for viral ecology and evolution

The comprehensive view of lispi-like virus diversity emerging from this data-mining effort reveals several key ecological and evolutionary patterns. First, extreme host specialization predominates despite substantial sampling, suggesting that molecular barriers to host range expansion are significant and that generalist viruses represent rare evolutionary achievements rather than common outcomes. Second, the viruses capable of generalist lifestyles show no consistent phylogenetic clustering, suggesting that host range expansion has evolved multiple times independently through convergent molecular adaptations.

Third, the strong correlation between agricultural importance/research intensity and virus detection highlights how anthropogenic factors shape our knowledge of viral diversity. Wild plant viruses remain vastly undersampled, representing an unknown risk for agricultural emergence and a substantial reservoir of unexplored biodiversity. Finally, the accelerating discovery rate indicates that plant virus diversity remains substantially underestimated even in well-studied systems. As sequencing costs continue declining and sampling extends into previously neglected plant groups and geographic regions, we anticipate continued discovery of novel viral lineages that will reshape our understanding of plant virome diversity, evolution, and ecology.

### 3.15 Strengths and limitations of sequence discovery through data mining

As previously demonstrated by Bejerman and colleagues [11, 42], independent validation through re-analyzing of SRA data can significantly enhance our understanding of RNA virus genomes with unique features. However, a key limitation of this approach is the lack of access to original biological material, which impedes replication and verification of the assembled viral genome sequences, an inherent weakness of data mining-based virus discovery. Additionally, potential issues such as contamination, low sequencing quality, sample spill-over, and other technical artifacts present a risk of generating false-positive, chimeric assemblies, or incorrect host assignments. Consequently, caution must be exercised when interpreting results derived from publicly available SRA data. To strengthen and validate our findings, we strongly recommend generating new RNAseq datasets from the predicted plant hosts. While our approach has inherent limitations, some aspects of our virus discovery strategy, such as iterative contig extension, and comprehensive sequence validation, can aid to alleviate some of these challenges and provide further evidence for identification. Nevertheless, it is key to emphasize that all virus-host associations and detections should be considered preliminary until confirmed through further investigation.

## 4. Conclusions

Our study reinforces and highlights the key importance of analyzing SRA public data as a valuable tool, not only for expediting the discovery of novel viruses but also for gaining insights into their evolutionary history and enhancing virus classification. Through this approach, we conducted a search for hidden lispi-like virus sequences, which significantly expanded the number of these viruses. Our analysis allowed us to shed light into its taxonomical classification, allocating a new plant-associated group of viruses to a new family within the *Mononegavirales* order; thus, being just the second family within this order that includes plant-associated members; which highlights their diversity and complex evolutionary dynamics of mononegaviruses. Some biological properties, such as transmission and viral particle morphology, were characterized for MSuV [28]; however, for all the other members, only the sequence was characterized. With the increasing reliance on sequence-based taxonomy, current classifications primarily reflect evolutionary relationships inferred from phylogenetic analyses of the L gene. In this context, the MSuV biological and sequence properties as well as the sequence properties of all newly identified viruses in our study, together with our phylogenetic analyses, support the assignment of these viruses within a new family, here proposed Masuviridae, within the order *Mononegavirales*.

## Acknowledgments

We would like to express genuine appreciation to the producers of the original sequence data used for this study, which are cited in **Table 1**. By ensuing open science practices with accessible raw sequence data in open public repositories, these authors supported contributions based on secondary data analyses.

## Author Contributions

Conceptualization, N.B and H.D; data analysis, N.B, D.H. and H.D; writing, original draft preparation, N.B and H.D. ; writing—review and editing, N.B, D.H, R. AC., D. QA. and H.D. All authors have read and agreed to the published version of the manuscript.

## Institutional Review Board Statement

Not applicable for studies not involving humans or animals.

## Informed Consent Statement

Not applicable for studies not involving humans.

## Data availability statement

Nucleotide sequence data reported are available in the Third Party Annotation Section of the DDBJ/ENA/GenBank databases under the accession numbers TPA: BK075274-BK075360 and can be found as **Supplementary Material 1** of this submission.

---

**Supp. Figure S1.** *Sequence-level characterisation of the L (P4) polymerase across 87 lispi-like viruses.* (A) Distribution of pairwise amino-acid identities, partitioned into within-host-family (blue) and between-host-family (grey) pairs (median 38.5%). (B) Pairwise identity matrix ordered by host botanical family (colour bar), showing coherent within-family blocks. (C) Per-site conservation (1 − H/H_max; 21-column sliding window) along the P4 alignment; the structurally resolved RdRp core (mean 0.69) is significantly more conserved than the structurally unresolved capping/methyltransferase/C-terminal region (mean 0.49; one-sided Mann–Whitney U, p ≈ 2 × 10⁻⁶⁵). Vertical marks indicate catalytic motifs A, B and C. (D) Sequence logos (information content, bits; n = 87) for motifs A, B and C. (E) Per-position percentage identity across the palmprint; catalytic residues (motif-A aspartate; motif-C G-D-N) in red. (F) Summary of catalytic-residue conservation: motif-A aspartate 100% (DFSKW invariant), motif-C G-D-N 100% (GDN in 87/87; no GDD or SDD variant), motif-B block GWEGQRQKGW 95.4% (83/87). The uniform GDN signature places P4 in the *Mononegavirales* / *Rhabdoviridae* RdRp catalytic class.

**Supp. Figure 2.** *Impact of Normalization on Virus Ranking* Comparison of virus prevalence rankings using raw library counts (A) versus normalized Virus Prevalence Score (VPS) (B). VPS is calculated as (Unique Hosts / √Libraries) × 100. Color gradients indicate detection frequency, with normalization revealing viruses that infect diverse hosts despite fewer total detections.

**Supp. Figure 3.** *Host Diversity versus Library Abundance* Scatter plot examining the relationship between total SRA libraries (x-axis, log scale) and number of unique host organisms (y-axis) for detected viruses. Bubble size represents normalized prevalence score (VPS), while color intensity indicates detection robustness (proportion of libraries where virus was detected).

**Supp. Figure 4.** *Library Redundancy Analysis* Comparison of total SRA libraries versus unique independent BioProjects for the top 30 viruses. Blue bars represent total library counts, while orange bars show unique BioProjects.

**Supp. Figure 5.** *Host Range Distribution: Monocots versus Dicots* Violin plots comparing the number of unique host organisms for dicot-dominated versus monocot-specific viruses.

**Supp. Figure 6.** *Tissue Tropism Patterns in Top 25 Viruses* Stacked bar chart showing proportional distribution of tissue types from which each virus was detected. Viruses are ordered by total detection frequency (left to right).

**Supp. Figure 7.** *Global Distribution of Viral Detections* World map showing 182 unique sampling locations across 252 countries. Bubble size indicates number of samples per location, while color represents virus richness (number of distinct viruses detected).

