## Supplementary material for "Data mining sheds light on a novel family of plant-associated negative-sense RNA viruses linked to lispiviruses": Table 1

**Table 1** . Summary of novel plant-associated lispi-like viruses identified from plant RNA-seq data available on NCBI.

| **Plant host** | **Taxa/**  **family** | **Virus name/**  **Abbreviation** | **Bioproject ID/**  **Data citation** | **Length (nt)/**  **coverage** | **Accession number** | **Protein ID** | **Length (aa)** | **Highest scoring virus- protein/*E*-value/query coverage%/identity% (Blast P)** |
| --- | --- | --- | --- | --- | --- | --- | --- | --- |
| floss flower  (*Ageratum houstonianum*) | Dicot/  *Asteraceae* | Ageratum lispi-like virus 1/  AgeLlV1 | PRJNA371565/  [86] | 13020/226.31X | BK075274 | N  P2  P3  L | 397  221  616  2359 | MSV-N/6e-22/52/31.1  no hits  no hits  MSV-L/0.0/100/34.31 |
| floss flower  (*Ageratum houstonianum*) | Dicot/  *Asteraceae* | Ageratum lispi-like virus 2/  AgeLlV2 | PRJNA371565/  [86] | 12086/218.45X | BK075275 | N  P2  P3  L | 384  236  644  2382 | MSV-N/3e-32/78/28.57  no hits  no hits  MSV-L/0.0/100/44.22 |
| Welsh onion  (*Allium fistulosum*) | Monocot/  *Allieae* | Allium lispi-like virus 1/  AllLlV1 | PRJNA542932/  [87] | 12047/44.05X | BK075276 | N  P2  P3  L | 379  218  676  2341 | MSV-N/4e-56/96/32.6  no hits  no hits  MSV-L/0.0/99/45.91 |
| wild garlic  (*Allium ursinum*) | Monocot/  *Allieae* | Allium lispi-like virus 2/  AllLlV2 | PRJNA542932/  [87] | 13318/421.95X | BK075277 | N  P2  P3  L | 417  271  695  2382 | MSV-N/8e-21/64/28.1  no hits  no hits  MSV-L/0.0/98/43.88 |
| alpine leek  (*Allium victoralis*) | Monocot/  *Allieae* | Allium lispi-like virus 3/  AllLlV3 | PRJNA633672/  [88] | 11800/163.99X | BK075278 | N  P2  P3  L | 363  219  627  2341 | MSV-N/8e-52/88/31.37  no hits  no hits  MSV-L/0.0/100/45.44 |
| common ragweed (*Ambrosia artemisiifolia*) | Dicot/  *Asteraceae* | Ambrosia lispi-like virus 1/  AmbLlV1 | PRJNA335689/  [89] | 12316/12.49X | BK075279 | N  P2  P3  L | 381  225  619  2374 | MSV-N/7e-39/71/34.8  no hits  no hits  MSV-L/0.0/100/41.85 |
| sand rock-cress  (*Arabidopsis arenosa*) | Dicot/  *Brassicaceae* | Arabidopsis lispi-like virus 1/  AraLlV1 | PRJNA575330/  [90] | 12590/23.01X | BK075280 | N  P2  P3  L | 383  243  614  2339 | MSV-N/2e-29/74/28.62  no hits  no hits  MSV-L/0.0/99/40.33 |
| dwarf mistletoe  (*Arceuthobium sichuanense*) | Dicot/  *Santalaceae* | Arceuthobium  lispi-like virus 1/  ArcLlV1 | PRJNA307530/  [91] | 12177/69.03X | BK075281 | N  P2  P3  L | 359  247  695  2353 | MSV-N/1e-24/74/31.7  no hits  no hits  MSV-L/0.0/100/40.19 |
| mountain tobacco  (*Arnica montana*) | Dicot/  *Asteraceae* | Arnica lispi-like virus 1/  ArnLlV1 | PRJNA233448/  [92] | 11732/23.84X | BK075282 | N  P2  P3  L | 367  220  655  2338 | MSV-N/7e-40/84/31.09  no hits  no hits  MSV-L/0.0/100/45.26 |
| Chinese milkvetch  (*Astragalus sinicus*) | Dicot/  *Fabaceae* | Astragalus lispi-like virus 1/  AstLlV1 | PRJNA690069/  [93] | 13281/29.38X | BK075283 | N  P2  P3  L | 370  239  669  2343 | MSV-N/2e-16/77/25.5  no hits  no hits  MSV-L/0.0/100/36.11 |
| Bai Zhu  (*Atractylodes macrocephala*) | Dicot/  *Asteraceae* | Atractylodes lispi-like virus 1/  AtrLlV1 | PRJNA419787/  [94] | 12540/56.48X | BK075284 | N  P2  P3  L | 406  211  636  2338 | MSV-N/2e-27/66/27.61  no hits  no hits  MSV-L/0.0/100/35.74 |
| black mustard  (*Brassica nigra*) | Dicot/  *Brassicaceae* | Brassica lispi-like virus 1/  BraLlV1 | PRJNA309775/  He, Z., York University, unpublished | 12516/16.43X | BK075285 | N  P2  P3  L | 405  244  564  2345 | MSV-N/9e-33/71/28.47  no hits  no hits  MSV-L/0.0/100/38.38 |
| rapeseed  (*Brassica napus*) | Dicot/  *Brassicaceae* | Brassica lispi-like virus 2_nap/  BraLlV2_nap | PRJNA549345/  [95] | 12489/38.49X | BK075286 | N  P2  P3  L | 402  244  573  2385 | MSV-N/5e-22/73/24.83  no hits  no hits  MSV-L/0.0/99/38.4 |
| turnip  (*Brassica rapa*) | Dicot/  *Brassicaceae* | Brassica lispi-like virus 2_rap/  BraLlV2_rap | PRJNA312975/  Zhong, L., SCAU, China, unpublished | 12253/15.22X | BK075287 | N  P2  P3  L | 402  244  573  2385 | MSV-N/5e-22/73/24.83  no hits  no hits  MSV-L/0.0/99/38.4 |
| short-pod mustard  (*Brassica Incana*) | Dicot/  *Brassicaceae* | Brassica lispi-like virus 3_inc/  BraLlV3_inc | PRJNA428769/  [96] | 12222/23.94X | BK075288 | N  P2  P3  L | 387  232  621  2341 | MSV-N/2e-34/87/30.68  no hits  no hits  MSV-L/0.0/99/38.78 |
| Indian mustard  (*Brassica insularis*) | Dicot/  *Brassicaceae* | Brassica lispi-like virus 3_ins/  BraLlV3_ins | PRJNA428769/  [96] | 12088/15.16X | BK075289 | N  P2  P3  L | 387  232  622  2341 | MSV-N/2e-34/87/30.68  no hits  no hits  MSV-L/0.0/99/38.78 |
| wild cabbage  (*Brassica oleracea*) | Dicot/  *Brassicaceae* | Brassica lispi-like virus 3_ole/  BraLlV3_ole | PRJNA428769/  [96] | 12183/42.49X | BK075290 | N  P2  P3  L | 387  232  621  2341 | MSV-N/2e-34/87/30.68  no hits  no hits  MSV-L/0.0/99/38.78 |
| mustard  (*Brassica rupestris*) | Dicot/  *Brassicaceae* | Brassica lispi-like virus 3_rup/  BraLlV3_rup | PRJNA428769/  [96] | 12230/14.47X | BK075291 | N  P2  P3  L | 387  232  621  2341 | MSV-N/2e-34/87/30.68  no hits  no hits  MSV-L/0.0/99/38.78 |
| Sicilian wild cabbage  (*Brassica villosa*) | Dicot/  *Brassicaceae* | Brassica lispi-like virus 3_vil/  BraLlV3_vil | PRJNA428769/  [96] | 12141/22.14X | BK075292 | N  P2  P3  L | 387  232  621  2341 | MSV-N/2e-34/87/30.68  no hits  no hits  MSV-L/0.0/99/38.78 |
| mantis orchid  (*Caladenia attigens*) | Monocot/  *Orchidaceae* | Caladenia lispi-like virus 1_att/  CalLlV1_att | PRJNA661963/  [97] | 11452/27.89X | BK075293 | N  P2  P3  L | 386  225  549  2324 | MSV-N/2e-34/78/30.59  no hits  no hits  MSV-L/0.0/100/40.11 |
| crab-lipped spider orchid  (*Caladenia plicata*) | Monocot/  *Orchidaceae* | Caladenia lispi-like virus 1_pli/  CalLlV1_pli | PRJNA384875/  [98] | 11404/21.43X | BK075294 | N  P2  P3  L | 386  225  549  2324 | MSV-N/2e-34/78/30.59  no hits  no hits  MSV-L/0.0/100/40.11 |
| Canberra spider orchid  (*Caladenia actensis*) | Monocot/  *Orchidaceae* | Caladenia lispi-like virus 2_act/  CalLlV2_act | PRJNA661963/  [97] | 11193/107.06X | BK075295 | N  P2  P3  L | 394  217  489  2314 | MSV-N/5e-37/76/32.57  no hits  no hits  MSV-L/0.0/100/39.86 |
| Arrowsmith spider orchid  (*Caladenia crebra*) | Monocot/  *Orchidaceae* | Caladenia lispi-like virus 2_cre/  CalLlV2_cre | PRJNA661963/  [97] | 11499/73.94X | BK075296 | N  P2  P3  L | 391  217  489  2314 | MSV-N/4e-33/74/32.23  no hits  no hits  MSV-L/0.0/100/39.86 |
| hemp  (*Cannabis sativa*) | Dicot/  *Cannabaceae* | Cannabis lispi-like virus 1/  CanLlV1 | PRJNA773629/  Yue, J., CAMS, China, unpublished | 13200/580.65X | BK075297 | N  P2  P3  L | 382  224  714  2344 | MSV-N/1e-38/80/32.79  no hits  no hits  MSV-L/0.0/100/42.93 |
| grand Shepherd's-purse  (*Capsella grandiflora*) | Dicot/  *Brassicaceae* | Capsella lispi-like virus 1/  CapLlV1 | PRJNA533007/  [99] | 12782/63.95X | BK075298 | N  P2  P3  L | 381  235  573  2344 | MSV-N/2e-22/90/26.42  no hits  no hits  MSV-L/0.0/100/39.68 |
| carrot  (*Daucus carota*) | Dicot/  Apiaceae | carrot lispi-like virus 1/  CarLlV1 | PRJNA574412/  [100] | 13434/524.80X | BK075299 | N  P2  P3  L | 458  220  644  2320 | MSV-N/4e-24/61/27.92  no hits  no hits  MSV-L/0.0/100/33.42 |
| pearl millet  (*Cenchrus americanus*) | Monocot/  *Poaceae* | Cenchrus lispi-like virus 1/  CenLlV1 | PRJNA625418/  Kumar, R., ICAR, India, unpublished | 12133/352.96X | BK075300 | N  P2  P3  L | 369  226  675  2328 | MSV-N/2e-38/91/27.84  no hits  no hits  MSV-L/0.0/100/44.8 |
| cornflower  (*Centaurea cyanus*) | Dicot/  *Asteraceae* | Centaurea lispi-like virus 1/ CentLlV1 | PRJNA577252/  [101] | 12996/12.07X | BK075301 | N  P2  P3  L | 393  215  636  2341 | MSV-N/4e-29/69/28.47  no hits  no hits  MSV-L/0.0/100/36.56 |
| common chicory  (*Cichorium intybus*) | Dicot/  *Asteraceae* | Cichorium lispi-like virus 1/  CicLlV1 | PRJNA328202/  [102] | 12530/18.57X | BK075302 | N  P2  P3  L | 389  217  644  2335 | MSV-N/8e-18/55/29.77  no hits  no hits  MSV-L/0.0/99/36.18 |
| desert ginseng  (*Cistanche deserticola*) | Dicot/  *Orobanchaceae* | Cistanche lispi-like virus 1/  CisLlV1 | PRJNA598927/  Li, M., Gansu, China, unpublished | 12415/237.02X | BK075303 | N  P2  P3  L | 396  245  659  2344 | MSV-N/3e-34/78/29.52  no hits  no hits  MSV-L/0.0/100/39-29 |
| desert ginseng  (*Cistanche deserticola*) | Dicot/  *Orobanchaceae* | Cistanche lispi-like virus 2/  CisLlV2 | PRJNA598927/  Li, M., Gansu, China, unpublished | 12552/112.82X | BK075304 | N  P2  P3  L | 396  245  659  2344 | MSV-N/5e-28/59/33.77  no hits  no hits  MSV-L/0.0/100/40.73 |
| golden thread  (*Coptis chinensis*) | Dicot/  *Ranunculaceae* | Coptis lispi-like virus 1/  CopLlV1 | PRJNA662860/  [103] | 12062/114.68X | BK075305 | N  P2  P3  L | 391  236  598  2342 | MSV-N/2e-46/86/32.25  no hits  no hits  MSV-L/0.0/100/40.3 |
| coriander  (*Coriandrum sativum*) | Dicot/  Apiaceae | Coriandrum lispi-like virus 1/  CorLlV1 | PRJNA531360/  [104] | 12694/14.25X | BK075306 | N  P2  P3  L | 431  215  691  2346 | MSV-N/1e-25/62/29.93  no hits  no hits  MSV-L/0.0/100/34.84 |
| coriander  (*Coriandrum sativum*) | Dicot/  Apiaceae | Coriandrum lispi-like virus 2/  CorLlV2 | PRJNA472685/  [105] | 12365/12.79X | BK075307 | N  P2  P3  L | 408  238  654  2343 | MSV-N/2e-38/86/33.25  no hits  no hits  MSV-L/0.0/100/44.68 |
| desert dodder  (*Cuscuta nevadensis*) | Dicot/  *Convolvulaceae* | Cuscuta lispi-like virus 1/  CusLlV1 | PRJNA561399/  Frangione E, U Toronto, Canada, unpublished | 12694/41.36X | BK075308 | N  P2  P3  L | 416  224  701  2329 | MSV-N/2e-29/65/29.89  no hits  no hits  MSV-L/0.0/100/35.25 |
| Taiwan spoon orchid  (*Cypripedium formosanum*) | Monocot/  *Orchidaceae* | Cypripedium lispi-like virus 1/  CypLlV1 | PRJNA277578/  [106] | 12935/71.96X | BK075309 | N  P2  P3  L | 369  226  684  2357 | MSV-N/3e-45/93/31.52  no hits  no hits  MSV-L/0.0/100/43.09 |
| Yellow slipper orchid  (*Cypripedium flavum*) | Monocot/  *Orchidaceae* | Cypripedium lispi-like virus 2/  CypLlV2 | PRJNA479379/  [107] | 12546/85.93X | BK075310 | N  P2  P3  L | 381  223  694  2342 | MSV-N/3e-34/76/30.03  no hits  no hits  MSV-L/0.0/100/39.95 |
| common spotted-orchid  (*Dactylorhiza fuchsii*) | Monocot/  *Orchidaceae* | Dactylorhiza lispi-like virus 1/  DacLlV1 | PRJNA317244/  [108] | 11802/18.81X | BK075311 | N  P2  P3  L | 368  223  628  2353 | MSV-N/2e-32/75/31.32  no hits  no hits  MSV-L/0.0/100/37.28 |
| common spotted-orchid  (*Dactylorhiza fuchsii*) | Monocot/  *Orchidaceae* | Dactylorhiza lispi-like virus 2/  DacLlV2 | PRJNA317244/  [108] | 12298/27.35X | BK075312 | N  P2  P3  L | 376  232  612  2402 | MSV-N/2e-36/79/31.79  no hits  no hits  MSV-L/0.0/100/35.05 |
| common spotted-orchid  (*Dactylorhiza fuchsii*) | Monocot/  *Orchidaceae* | Dactylorhiza lispi-like virus 3/  DacLlV3 | PRJNA317244/  [108] | 11960/50.36X | BK075313 | N  P2  P3  L | 367  222  620  2353 | MSV-N/6e-32/77/31.27  no hits  no hits  MSV-L/0.0/100/37.33 |
| Venus flytrap  (*Dionaea muscipula*) | Dicot/  Droseraceae | Dionaea lispi-like virus 1/  DioLlV1 | PRJNA203407/  [109] | 12600/266.77X | BK075314 | N  P2  P3  L | 369  236  658  2344 | MSV-N/2e-43/75/34.04  no hits  no hits  MSV-L/0.0/100/41.73 |
| violet helleborine  (*Epipactis purpurata*) | Monocot/  *Orchidaceae* | Epipactis lispi-like virus 1/  EpiLlV1 | PRJNA450088/  [110] | 13620/169.66X | BK075315 | N  P2  P3  L | 374  219  651  2345 | MSV-N/1e-46/79/34.01  no hits  no hits  MSV-L/0.0/100/41.05 |
| violet helleborine  (*Epipactis purpurata*) | Monocot/  *Orchidaceae* | Epipactis lispi-like virus 2/  EpiLlV2 | PRJNA450088/  [110] | 12946/120.39X | BK075316 | N  P2  P3  L | 391  219  678  2349 | MSV-N/4e-37/85/29.55  no hits  no hits  MSV-L/0.0/100/40.21 |
| shortscape fleabane  (*Erigeron breviscapus*) | Dicot/  *Asteraceae* | Erigeron lispi-like virus 1/  EriLlV1 | PRJNA293262/  [111] | 11755/33.58X | BK075317 | N  P2  P3  L | 375  222  649  2338 | MSV-N3e-43/95/31.97  no hits  no hits  MSV-L/0.0/100/45.88 |
| garden rocket  (*Eruca vesicaria*) | Dicot/  *Brassicaceae* | Eruca lispi-like virus 1_eru/  EruLlV1_eru | PRJNA475309/  [112] | 12152/53.71X | BK075318 | N  P2  P3  L | 401  243  569  2342 | MSV-N/2e-23/72/26.71  no hits  no hits  MSV-L/0.0/99/38.5 |
| wallflower  (*Erysimum cheiri*) | Dicot/  *Brassicaceae* | Eruca lispi-like virus 1_ery/  EruLlV1_ery | PRJNA362874/  [113] | 12427/73.70X | BK075319 | N  P2  P3  L | 401  243  569  2395 | MSV-N/2e-23/72/26.71  no hits  no hits  MSV-L/0.0/99/37.72 |
| Bastetan wallflower  (*Erysimum bastetanum*) | Dicot/  *Brassicaceae* | Erysimum lispi-like virus 1/  EryLlV1 | PRJNA607615/  [114] | 12670/275.88X | BK075320 | N  P2  P3  L | 411  235  587  2389 | MSV-N/6e-19/74/25.9  no hits  no hits  MSV-L/0.0/98/37.17 |
| niger  (*Guizotia abyssinica*) | Dicot/  *Asteraceae* | Guizotia lispi-like virus 1/  GuiLlV1 | PRJNA763316/  [115] | 12071/23.56X | BK075321 | N  P2  P3  L | 377  221  717  2336 | MSV-N/2e-42/88/30.27  no hits  no hits  MSV-L/0.0/100/45.42 |
| marsh-fragrant orchid  (*Gymnadenia densiflora*) | Monocot/  *Orchidaceae* | Gymnadenia lispi-like virus 1/  GymLlV1 | PRJNA504609/  [116] | 12203/145.36X | BK075322 | N  P2  P3  L | 367  222  634  2355 | MSV-N/1e-32/76/31.01  no hits  no hits  MSV-L/// |
| very small catchfly  (*Heliosperma pusillum*) | Dicot/  *Caryophyllaceae* | Heliosperma lispi-like virus 1/  HelLlV1 | PRJNA760819/  [117] | 12884/271.81X | BK075323 | N  P2  P3  L | 396  233  632  2362 | MSV-N/6e-21/85/26.53  no hits  no hits  MSV-L/0.0/99/36.89 |
| wild hogweed  (*Heracleum moellendorffii*) | Dicot/  *Apiaceae* | Heracleum lispi-like virus 1/  HerLlV1 | PRJNA438242  [118] | 12352/16.98X | BK075324 | N  P2  P3  L | 402  237  654  2342 | MSV-N/9e-31/88/27.81  no hits  no hits  MSV-L/0.0/100/44.28 |
| lily  (*Ixiolirion sp*.) | Monocot/  *Ixioliriaceae* | Ixiolirion lispiu-like virus 1/  IxiLlV1 | PRJNA412930/  [119] | 11807/80.61X | BK075325 | N  P2  P3  L | 385  247  583  2341 | MSV-N/4e-29/90/29.05  no hits  no hits  MSV-L/0.0/99/39.92 |
| Indian lettuce  (*Lactuca indica*) | Dicot/  *Asteraceae* | Lactuca lispi-like virus 1/  LacLlV1 | PRJNA386610/  [120] | 12952/52.91X | BK075326 | N  P2  P3  L | 388  223  632  2345 | MSV-N/2e-24/85/24.77  no hits  no hits  MSV-L/0.0/98/39.84 |
| sea lavender  (*Limonium bicolor*) | Dicot/  *Plumbaginaceae* | Limonium lispi-like virus 1/  LimLlV1 | PRJNA752802/  [121] | 12676/14.84X | BK075327 | N  P2  P3  L | 377  220  630  2340 | MSV-N/1e-31/78/31.65  no hits  no hits  MSV-L/0.0/100/38.93 |
| anual mercuri  (*Merculiaris annua*) | Dicot/  *Euphorbiaceae* | Mercurialis lispi-like virus 1/  MerLlV1 | PRJNA369310/  [122] | 13024/174.77X | BK075328 | N  P2  P3  L | 412  236  615  2325 | MSV-N/7e-20/62/29.17  no hits  no hits  MSV-L/0.0/98/37.02 |
| ghost plant  (*Monotropastrum humile*) | Dicot/  *Ericaceae* | Monotropastrum lispi-like virus 1/  MonLlV1 | PRJDB3033/  NIBB, Japan,  unpublished | 12029/101.32X | BK075329 | N  P2  P3  L | 375  227  632  2347 | MSV-N/4e-35/74/33.09  no hits  no hits  MSV-L/0.0/100/40.92 |
| violet cabbage  (*Moricandia arvensis*) | Dicot/  *Brassicaceae* | Moricandia lispi-like virus 1/  MorLlV1 | PRJNA604514/  [123] | 12409/228.54X | BK075330 | N  P2  P3  L | 402  244  571  2346 | MSV-N/4e-21/69/24.55  no hits  no hits  MSV-L/0.0/98/38.19 |
| dark bee orchid  (*Ophrys fusca*) | Monocot/  *Orchidaceae* | Ophrys lispi-like virus 1/  OphLlV1 | PRJNA574279  [116] | 12020/15.87X | BK075331 | N  P2  P3  L | 371  231  637  2353 | MSV-N/4e-31/90/29.13  no hits  no hits  MSV-L/0.0/100/37.95 |
| early spider orchid  (*Ophrys sphegodes*) | Monocot/  *Orchidaceae* | Ophrys lispi-like virus 2/  OphLlV2 | PRJNA574279  [116] | 11574/34.56X | BK075332 | N  P2  P3  L | 371  223  638  2353 | MSV-N/2e-32/80/29.34  no hits  no hits  MSV-L/0.0/100/37.45 |
| Chinese violet cree  (*Orychophragmus violaceus*) | Dicot/  *Brassicaceae* | Orychophragmus lispi-like virus 1/  OryLlV1 | PRJNA431031/  Zhong, L., Sichuan U, China, unpublished | 12197/39.46X | BK075333 | N  P2  P3  L | 390  231  597  2340 | MSV-N/4e-28/84/28.57  no hits  no hits  MSV-L/0.0/93/41.09 |
| herb Paris  (*Paris polyphylla*) | Monocot/  *Melanthiaceae* | Paris lispi-like virus 1/  ParLlV1 | PRJNA783190/  [124] | 12001/85.56X | BK075334 | N  P2  P3  L | 390  226  660  2348 | MSV-N/2e-25/76/29.57  no hits  no hits  MSV-L/0.0/99/36.16 |
| desertbells  (*Phacelia campanularia*) | Dicot/  *Boraginaceae* | Phacelia lispi-like virus 1/ PhaLlV1 | PRJEB21674/  OneKP project, unpublished | 12932/22.04X | BK075335 | N  P2  P3  L | 374  234  618  2343 | MSV-N/1e-30/75/29.79  no hits  no hits  MSV-L/0.0/100/36.25 |
| ribwort plantain  (*Plantago lanceolata*) | Dicot/  *Plantaginaceae* | Plantago lispi-like virus 1/  PlaLlV1 | PRJNA636383/  [125] | 12367/32.36X | BK075336 | N  P2  P3  L | 465  223  587  2340 | MSV-N/5e-13/44/29.14  no hits  no hits  MSV-L/0.0/99/35.68 |
| garden radish  (*Raphanus sativus*) | Dicot/  *Brassicaceae* | Raphanus lispi-like virus 1/  RapLlV1 | PRJNA650223/  [126] | 12500/515.06X | BK075337 | N  P2  P3  L | 388  239  597  2340 | MSV-N/1e-18/88/24.93  no hits  no hits  MSV-L/0.0/99/39.37 |
| stinkhorn clubhead  (*Rhopalocnemis phalloides*) | Dicot/  *Balanophoraceae* | Rhopalocnemis lispi-like virus 1/  RhoLlV1 | PRJNA737177/  [127] | 11830/129.19X | BK075338 | N  P2  P3  L | 396  259  561  2346 | MSV-N/5e-22/56/30.97  no hits  no hits  MSV-L/0.0/100/37.83 |
| stinkhorn clubhead  (*Rhopalocnemis phalloides*) | Dicot/  *Balanophoraceae* | Rhopalocnemis lispi-like virus 2/  RhoLlV2 | PRJNA737177/  [127] | 12460/183.34X | BK075339 | N  P2  P3  L | 381  254  568  2343 | MSV-N/1e-37/87/30.54  no hits  no hits  MSV-L/0.0/100/38.65 |
| stinkhorn clubhead  (*Rhopalocnemis phalloides*) | Dicot/  *Balanophoraceae* | Rhopalocnemis lispi-like virus 3/  RhoLlV3 | PRJNA737177/  [127] | 11760/132.53X | BK075340 | N  P2  P3  L | 395  234  545  2344 | MSV-N/1e-36/70/31.05  no hits  no hits  MSV-L/0.0/100/38.26 |
| stinkhorn clubhead  (*Rhopalocnemis phalloides*) | Dicot/  *Balanophoraceae* | Rhopalocnemis lispi-like virus 4/  RhoLlV4 | PRJNA495456/  [128] | 12177/280.79X | BK075341 | N  P2  P3  L | 401  240  560  2340 | MSV-N/7e-37/81/31.63  no hits  no hits  MSV-L/0.0/100/37.76 |
| stinkhorn clubhead  (*Rhopalocnemis phalloides*) | Dicot/  *Balanophoraceae* | Rhopalocnemis lispi-like virus 5/  RhoLlV5 | PRJNA495456/  [128] | 12030/681.54X | BK075342 | N  P2  P3  L | 401  234  546  2344 | MSV-N/1e-29/52/32.31  no hits  no hits  MSV-L/0.0/100/38.46 |
| stinkhorn clubhead  (*Rhopalocnemis phalloides*) | Dicot/  *Balanophoraceae* | Rhopalocnemis lispi-like virus 6/  RhoLlV6 | PRJNA495456/  [128] | 11897/909.17X | BK075343 | N  P2  P3  L | 405  234  546  2344 | MSV-N/1e-29/51/33.97  no hits  no hits  MSV-L/0.0/100/38.08 |
| stinkhorn clubhead  (*Rhopalocnemis phalloides*) | Dicot/  *Balanophoraceae* | Rhopalocnemis lispi-like virus 7/  RhoLlV7 | PRJNA495456/  [128] | 11955/561.56X | BK075344 | N  P2  P3  L | 387  250  571  2345 | MSV-N/3e-26/72/27.96  no hits  no hits  MSV-L/0.0/100/37.92 |
| heartwing dock  (*Rumex hastatulus*) | Dicot/  *Polygonaceae* | Rumex lispi-like virus 1/  RumLlV1 | PRJNA472058/  [129] | 12245/277.42X | BK075345 | N  P2  P3  L | 378  223  652  2340 | MSV-N/9e-50/76/36.68  no hits  no hits  MSV-L/0.0/100/45.74 |
| alpine sheep sorrell  (*Rumex paucifolius*) | Dicot/  *Polygonaceae* | Rumex lispi-like virus 2/  RumLlV2 | PRJNA698922/  [130] | 11965/91.58X | BK075346 | N  P2  P3  L | 378  222  656  2340 | MSV-N/5e-45/76/34.59  no hits  no hits  MSV-L/0.0/100/45.4 |
| Chinese laserwort  (*Saposhnikovia divaricata*) | Dicot/  *Apiaceae* | Saposhnikovia lispi-like virus 1/  SapLlV1 | PRJNA613937/  [131] | 13007/12.98X | BK075347 | N  P2  P3  L | 467  220  665  2392 | MSV-N/4e-26/59/29.35  no hits  no hits  MSV-L/0.0/99/34.33 |
| pincushion flower  (*Scabiosa columbaria*) | Dicot/  *Caprifoliaceae* | Scabiosa lispi-like virus 1/  ScaLlV1 | PRJNA80113/  [132] | 12732/82.29X | BK075348 | N  P2  P3  L | 403  228  696  2345 | MSV-N/9e-22/87/23.51  no hits  no hits  MSV-L/0.0/100/43.17 |
| white campion  (*Silene latifolia*) | Dicot/  *Caryophyllaceae* | Silene lispi-like virus 1_lat/  SilLlV1_lat | PRJEB39526/  [133] | 12359/114.54X | BK075349 | N  P2  P3  L | 388  263  621  2334 | MSV-N/2e-18/72/27.58  no hits  no hits  MSV-L/0.0/100/39.64 |
| Hueffelii catchfly  (*Silene heuffelii*) | Dicot/  *Caryophyllaceae* | Silene lispi-like virus 1_heu/  SilLlV1_heu | PRJEB39526/  [133] | 12347/128.39X | BK075350 | N  P2  P3  L | 388  263  621  2334 | MSV-N/2e-18/72/27.58  no hits  no hits  MSV-L/0.0/100/39.64 |
| bladder campion  (*Silene vulgaris*) | Dicot/  *Caryophyllaceae* | Silene lispi-like virus 2/  SilLlV2 | PRJEB39526/  [133] | 12174/60.44X | BK075351 | N  P2  P3  L | 390  250  626  2334 | MSV-N/2e-21/73/28.32  no hits  no hits  MSV-L/0.0/99/38.93 |
| Limpricht's Sinalliaria  (*Sinalliaria limprichtiana*) | Dicot/  *Brassicacea* | Sinalliaria lispi-like virus 1/  SinLlV1 | PRJNA428441/  [134] | 12804/45.07X | BK075352 | N  P2  P3  L | 407  232  587  2375 | MSV-N/7e-19/6/25.09  no hits  no hits  MSV-L/0.0/100/37.44 |
| wild mustard  (*Sinapis arvensis*) | Dicot/  *Brassicacea* | Sinapis lispi-like virus 1/  SinaLlV1 | PRJNA232677/  [135] | 12112/61.46X | BK075353 | N  P2  P3  L | 385  233  601  2342 | MSV-N/1e-24/86/26.39  no hits  no hits  MSV-L/0.0/99/39.88 |
| Katherine sorghum  (*Sorghum macrospermum*) | Monocot/  *Poaceae* | Sorghum lispi-like virus 1/  SorLlV1 | PRJNA736757/  [136] | 12155/434.64X | BK075354 | N  P2  P3  L | 374  230  623  2335 | MSV-N/8e-43/83/29.94  no hits  no hits  MSV-L/0.0/100/41.46 |
| narrow-saline sipweed  (*Suaeda salsa*) | Dicot/  *Amaranthaceae* | Suaeda lispi-like virus 1/  SuaLlV1 | PRJNA383092/  Zheng, M., FIO, China, unpublished | 12704/13.26X | BK075355 | N  P2  P3  L | 390  239  699  2365 | MSV-N/2e-31/69/31.34  no hits  no hits  MSV-L/0.0/98/36.93 |
| sunflower  (*Helianthus annuus*) | Dicot/  *Asteraceae* | sunflower lispi-like virus 1/  SunLlV1 | PRJNA408292/  [137] | 13579/19.07X | BK075356 | N  P2  P3  L | 393  215  599  2341 | MSV-N/1e-13/54/25.57  no hits  no hits  MSV-L/0.0/100/34.38 |
| Dalmatian chrysanthemum  (*Tanacetum cinerariifolium*) | Dicot/  *Asteraceae* | Tanacetum lispi-like virus 1/  TanLlV1 | PRJNA399494/  [138] | 11778/12.27X | BK075357 | N  P2  P3  L | 368  221  663  2338 | MSV-N/4e-37/75/30.58  no hits  no hits  MSV-L/0.0/100/45.14 |
| bull clover  (*Trioflium fucatum*) | Dicot/  *Fabaceae* | Trifolium lispi-like virus 1/  TriLlV1 | PRJNA311722/  [139] | 12046/97.71X | BK075358 | N  P2  P3  L | 374  225  609  2391 | MSV-N/5e-41/75/30.36  no hits  no hits  MSV-L/0.0/98/45.49 |
| broad bean  (*Vicia faba*) | Dicot/  *Fabaceae* | Vicia lispi-like virus 1/  VicLlV1 | PRJNA679419/  [140] | 13583/122.89X | BK075359 | N  P2  P3  L | 431  253  661  2342 | MSV-N/2e-14/56/26.72  no hits  no hits  MSV-L/0.0/100/36.32 |
| Korean mistletoe  (*Viscum coloratum*) | Dicot/  *Santalaceae* | Viscum lispi-like virus 1/  VisLlV1 | PRJNA350768/  [141] | 11348/28.59X | BK075360 | N  P2  P3  L | 370  219  606  2343 | MSV-N/1e-31/86/28.83  no hits  no hits  MSV-L/0.0/100/39.26 |
