## Supplementary material for "Data mining sheds light on a novel family of plant-associated negative-sense RNA viruses linked to lispiviruses": Table 2

**Table 2**. Consensus conserved lispi-like viruses gene junction sequences

| **Virus*** | **3´end mRNA** | **intergenic spacer** | **5´end mRNA** |
| --- | --- | --- | --- |
| MSuV | AUCUUUUUU | CAC | UCU |
| AgeLlV1 | AAUAUUUUU | CAC | UUG |
| AgeLlV2 | UUAUUUUUU | CAC | UCC |
| AllLlV1 | AUCUUUUUU | CAC | UCC |
| AllLlV2 | AACUUUUUU | CAC | UUU |
| AllLlV3 | AUCUUUUUU | CAC | UGG |
| AmbLlV1 | AAAUUUUUU | CAC | UUU |
| AraLlV1 | AUUCUUUUU | CGC | UCU |
| ArcLlV1 | AUCUUUUUU | CAU | UCU |
| ArnLlV1 | AUCUUUUUU | CAC | UCU |
| AstLlV1 | AAAUUUUUU | CAC | UCU |
| AtrLlV1 | UUAUUUUUU | CAC | UCU |
| BraLlV1 | AUAUUUUUU | CUC | UUU |
| BraLlV2 | UAAUUUUUU | CAC | UUG |
| BraLlV3 | AAUCUUUUU | CGC | UCC |
| CalLlV1 | AUCUUUUUU | CAC | UCC |
| CalLlV2 | AAUCUUUUU | CUC | UCU |
| CanLlV1 | AUCUUUUUU | CAC | UCC |
| CapLlV1 | AUUCUUUUU | CGC | UUC |
| CarLlV1 | AAAUUUUUU | CAC | UCG |
| CenLlV1 | AUAUUUUUU | CAC | UCC |
| CentLlV1 | AUAUUUUUU | CAC | UCG |
| CicLlV1 | AUUAUUUUU | CAC | UUU |
| CisLlV1 | AUCUUUUUU | CGC | UCU |
| CisLlV2 | UACUUUUUU | CGC | UUG |
| CopLlV1 | AUCUUUUUU | CAC | UCC |
| CorLlV1 | AAAUUUUUU | CGC | UUU |
| CorLlV2 | AUCUUUUUU | CAC | UCU |
| CusLlV1 | UAAUUUUUU | CAC | UCU |
| CypLlV1 | CGUUUUUUU | CGC | UCU |
| CypLlV2 | AUGUUUUUU | CAC | UCC |
| DacLlV1 | AACUUUUUU | CAC | UCU |
| DacLlV2 | AUCUUUUUU | CGC | UCC |
| DacLlV3 | AUCUUUUUU | CAC | UCU |
| DioLlV1 | UUGUUUUUU | CAC | UCC |
| EpiLlV1 | UUCUUUUUU | CAC | UCU |
| EpiLlV2 | UAAUUUUUU | CGC | UCC |
| EriLlV1 | UUAUUUUUU | CAC | UCC |
| EruLlV1 | AUAUUUUUU | CAC | UUA |
| EryLlV1 | AUAUUUUUU | CGC | UUA |
| GuiLlV1 | AUCUUUUUU | CGC | UCC |
| GymLlV1 | AUCUUUUUU | CAC | UCU |
| HelLlV1 | AUAUUUUUU | CAU | UCA |
| HerLlV1 | AUCUUUUUU | CAC | UCU |
| IxiLlV1 | AUUCUUUUU | CGC | UCU |
| LacLlV1 | AUCUUUUUU | CAC | UCG |
| LimLlV1 | AUAUUUUUU | CAC | UUC |
| MerLlV1 | AUAUUUUUU | CAC | UCA |
| MonLlV1 | AUCUUUUUU | CGC | UCU |
| MorLlV1 | AUAUUUUUU | CAC | UCU |
| OphLlV1 | AUUCUUUUU | CAC | UCU |
| OphLlV2 | AUUCUUUUU | CAC | UCU |
| OryLlV1 | AUUCUUUUU | CAC | UCC |
| ParLlV1 | AUCUUUUUU | CAC | UCC |
| PhaLlV1 | AUGUUUUUU | CAC | UCU |
| PlaLlV1 | AAAUUUUUU | CAU | UCU |
| RapLlV1 | AUUCUUUUU | CGC | UCC |
| RhoLlV1 | AUCUUUUUU | CGC | UCU |
| RhoLlV2 | AUCUUUUUU | CGC | UCU |
| RhoLlV3 | UUGUUUUUU | CGC | UCU |
| RhoLlV4 | UUAUUUUUU | CGC | UCU |
| RhoLlV5 | AUAUUUUUU | CGC | UCU |
| RhoLlV6 | AUAUUUUUU | CAC | UCU |
| RhoLlV7 | AUCUUUUUU | CGC | UCU |
| RumLlV1 | AUAUUUUUU | CGC | UCU |
| RumLlV2 | AACUUUUUU | CAC | UCU |
| SapLlV1 | AUAUUUUUU | CAC | UCU |
| ScaLlV1 | AUGUUUUUU | CAC | UCU |
| SilLlV1 | AUUCUUUUU | CGC | UCU |
| SilLlV2 | AUUCUUUUU | CGC | UCU |
| SinLlV1 | UUCUUUUUU | CGC | UCU |
| SinaLlV1 | AUUCUUUUU | CAC | UCC |
| SorLlV1 | AUCUUUUUU | CAC | UCC |
| SuaLlV1 | AUCUUUUUU | CGC | UCC |
| SunLlV1 | AUAAUUUUU | CUC | UCG |
| TanLlV1 | AUCUUUUUU | CAC | UCC |
| TriLlV1 | AUCUUUUUU | CAC | UCC |
| VicLlV1 | UAAUUUUUU | CAC | UCG |
| VisLlV1 | AUUCUUUUU | CAC | UCU |

The consensus gene junction sequences of the viruses identified in this study are highlighted in light grey. * Names and abbreviations of newly identified viruses are listed in Table 1; MSuV stands for maize suscal virus.
