## Supplementary figures and images for "Data mining sheds light on a novel family of plant-associated negative-sense RNA viruses linked to lispiviruses"

### Supp. Fig. 1

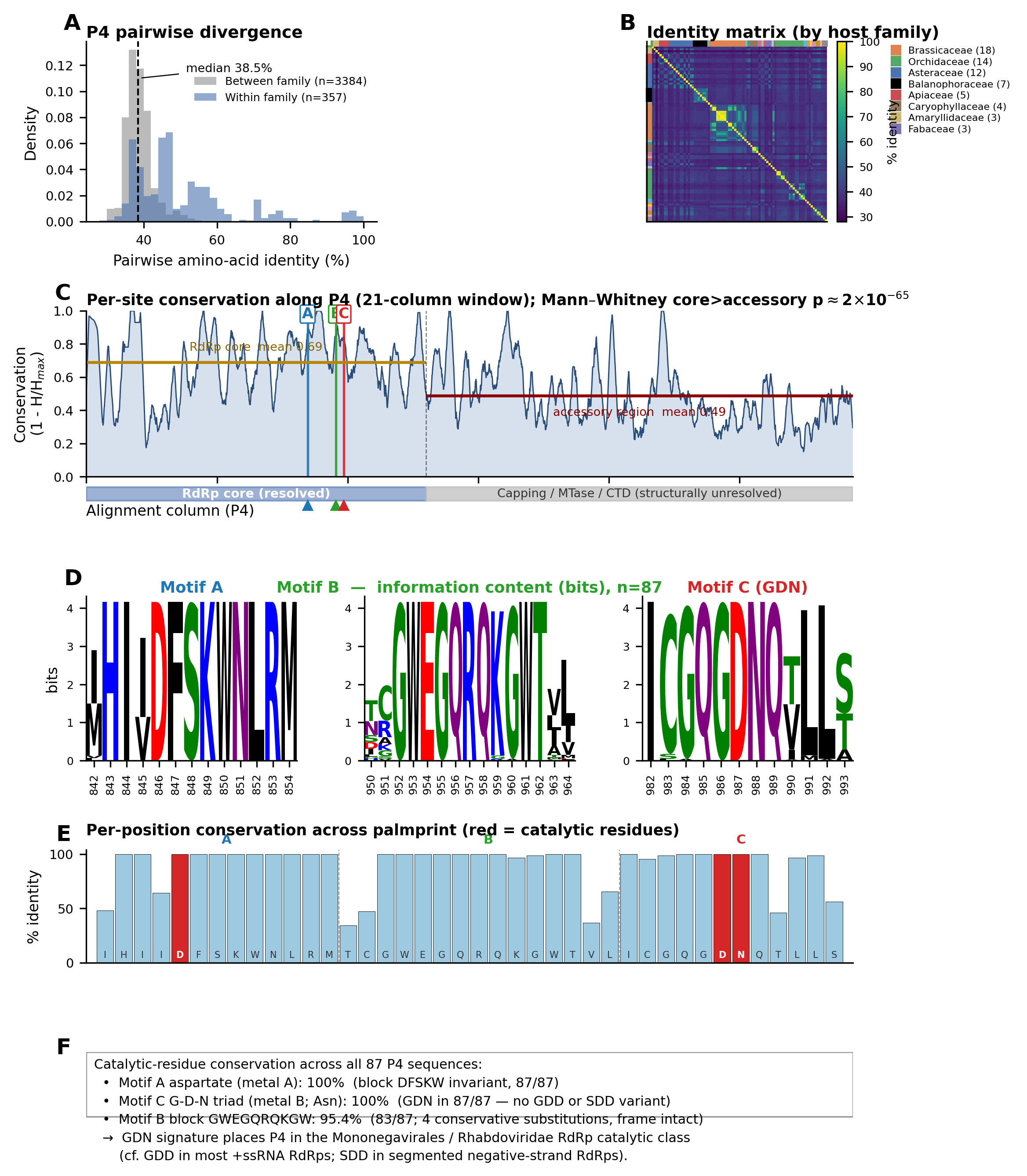

### Supp. Fig. 2

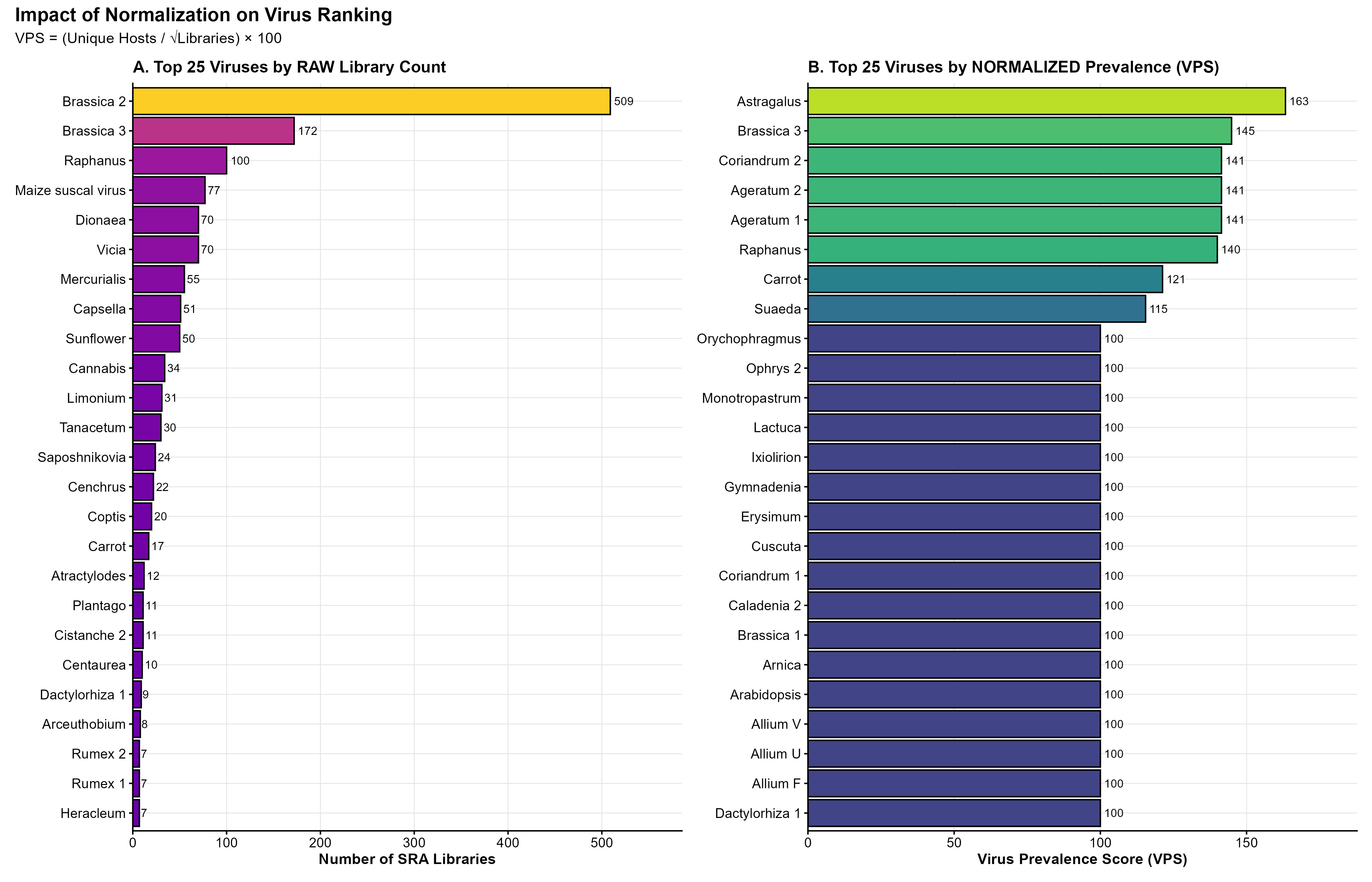

### Supp. Fig. 3

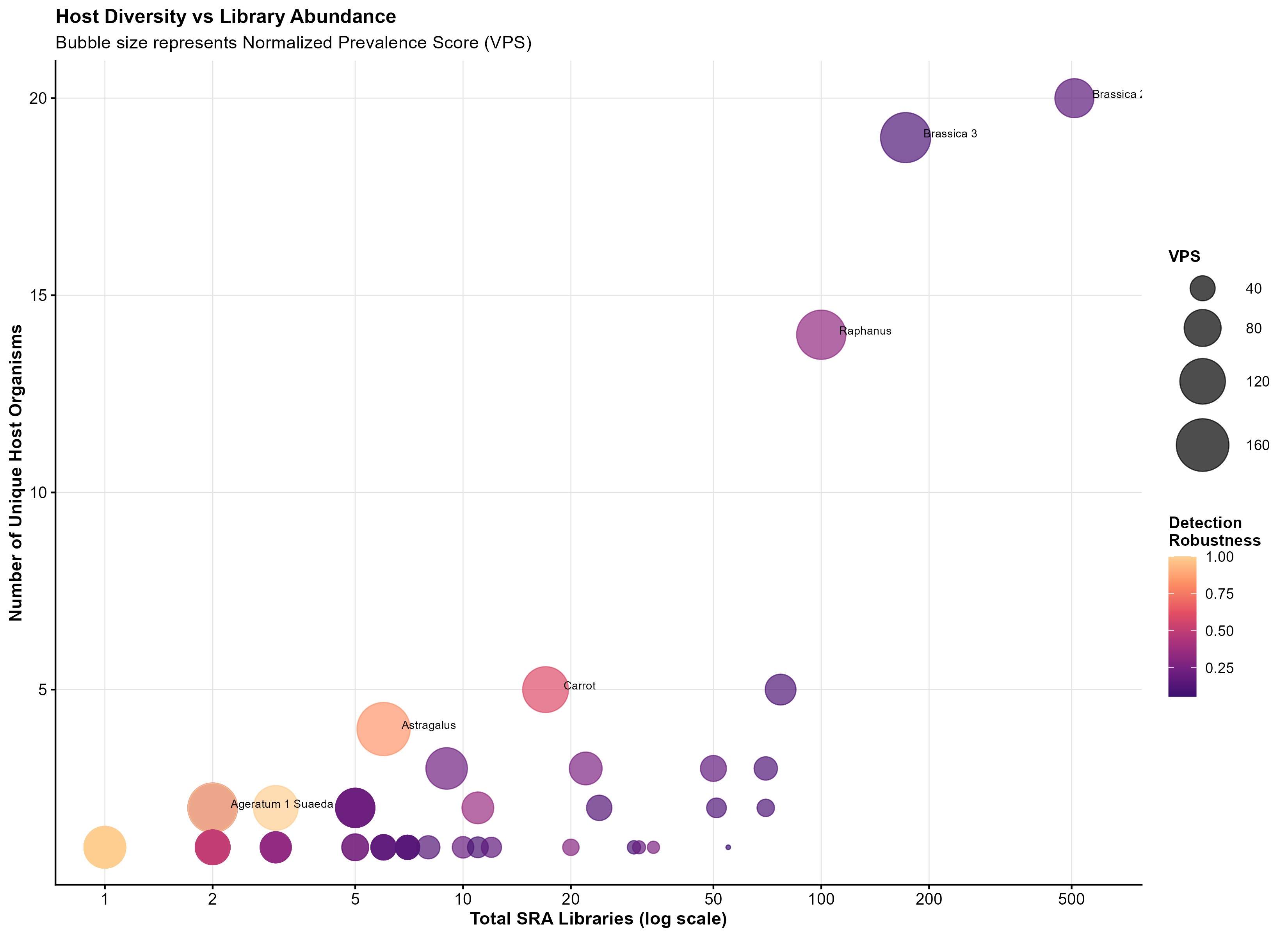

### Supp. Fig. 4

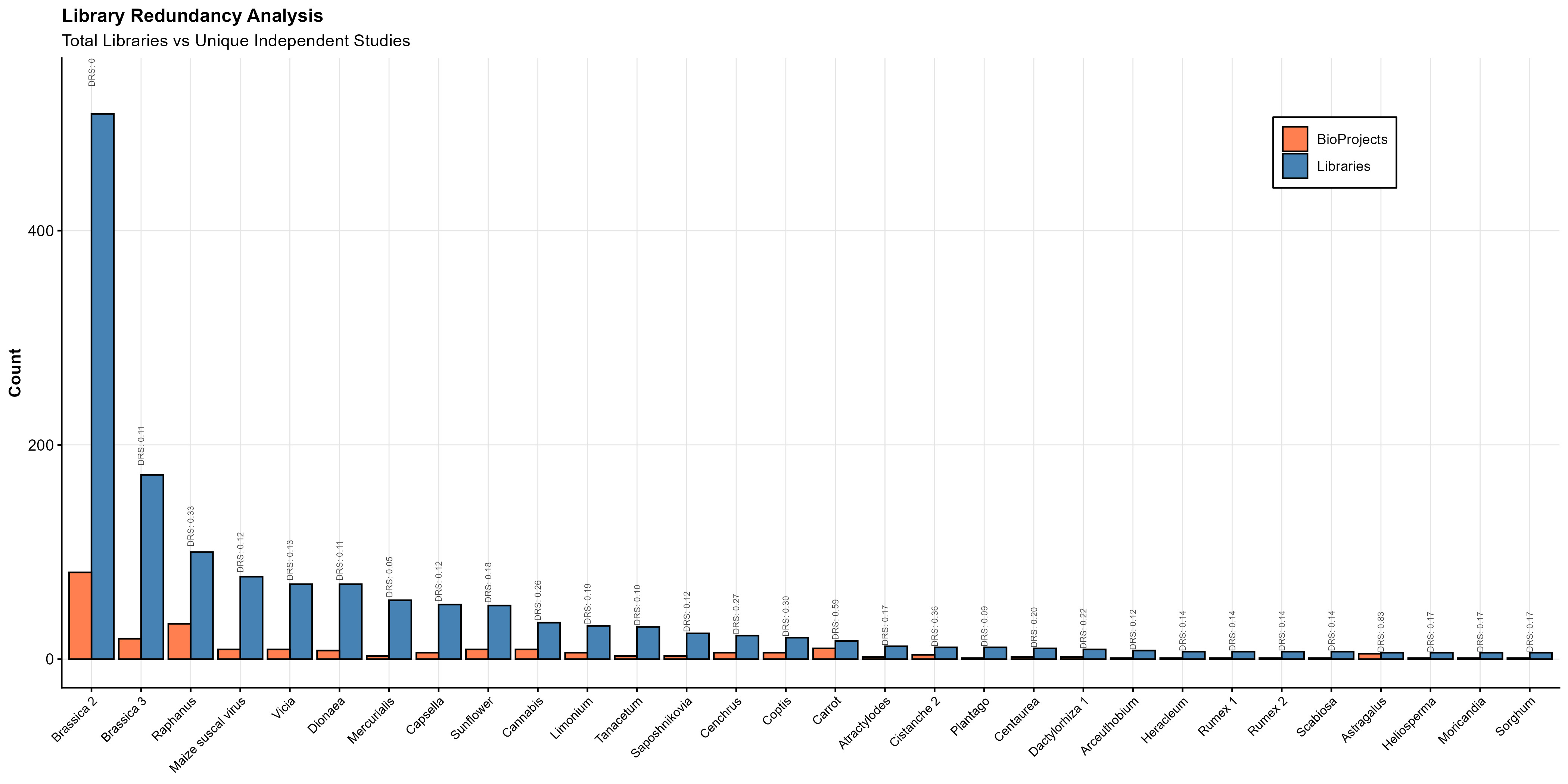

### Supp. Fig. 5

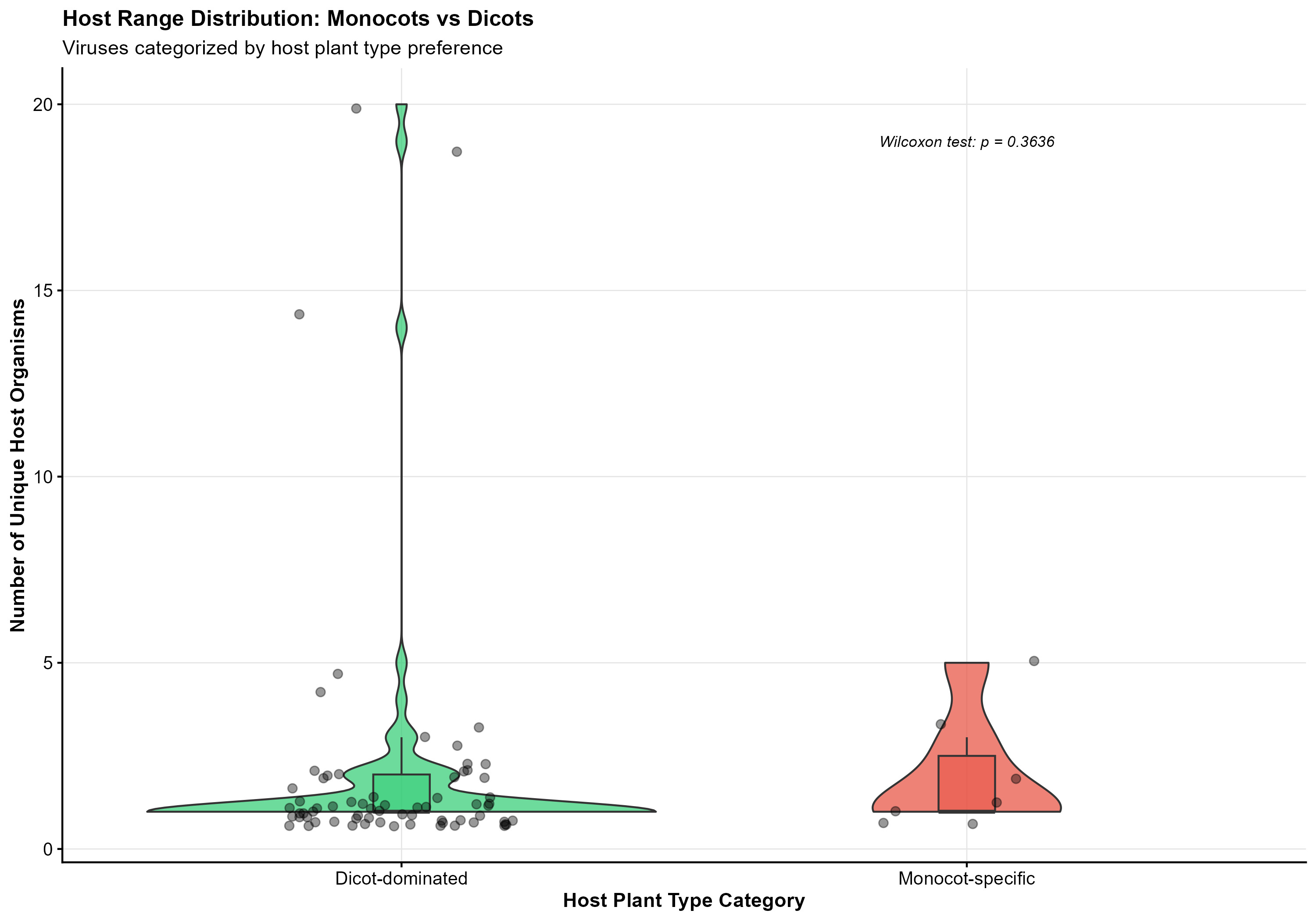

### Supp. Fig. 6

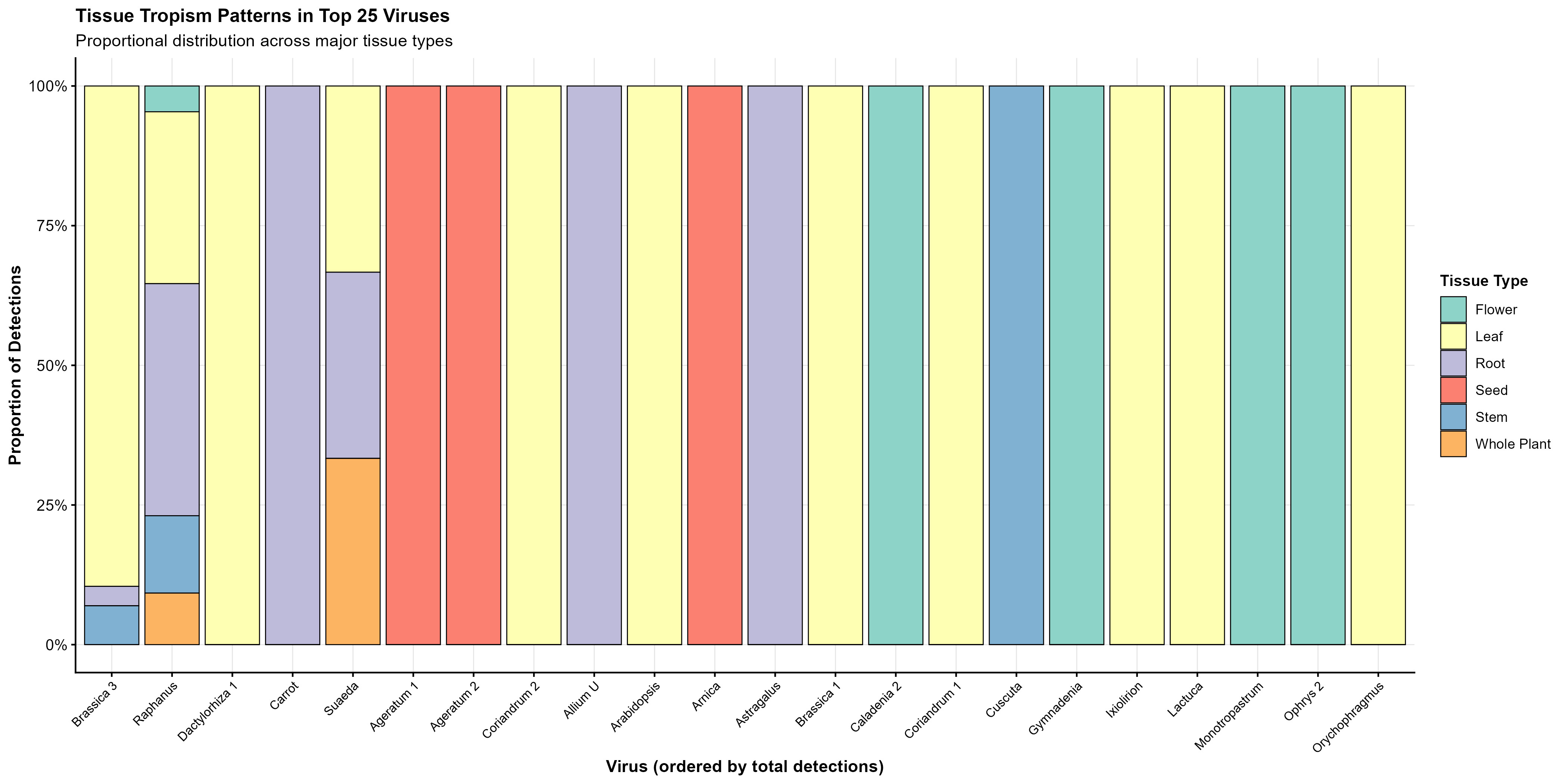

### Supp. Fig. 7

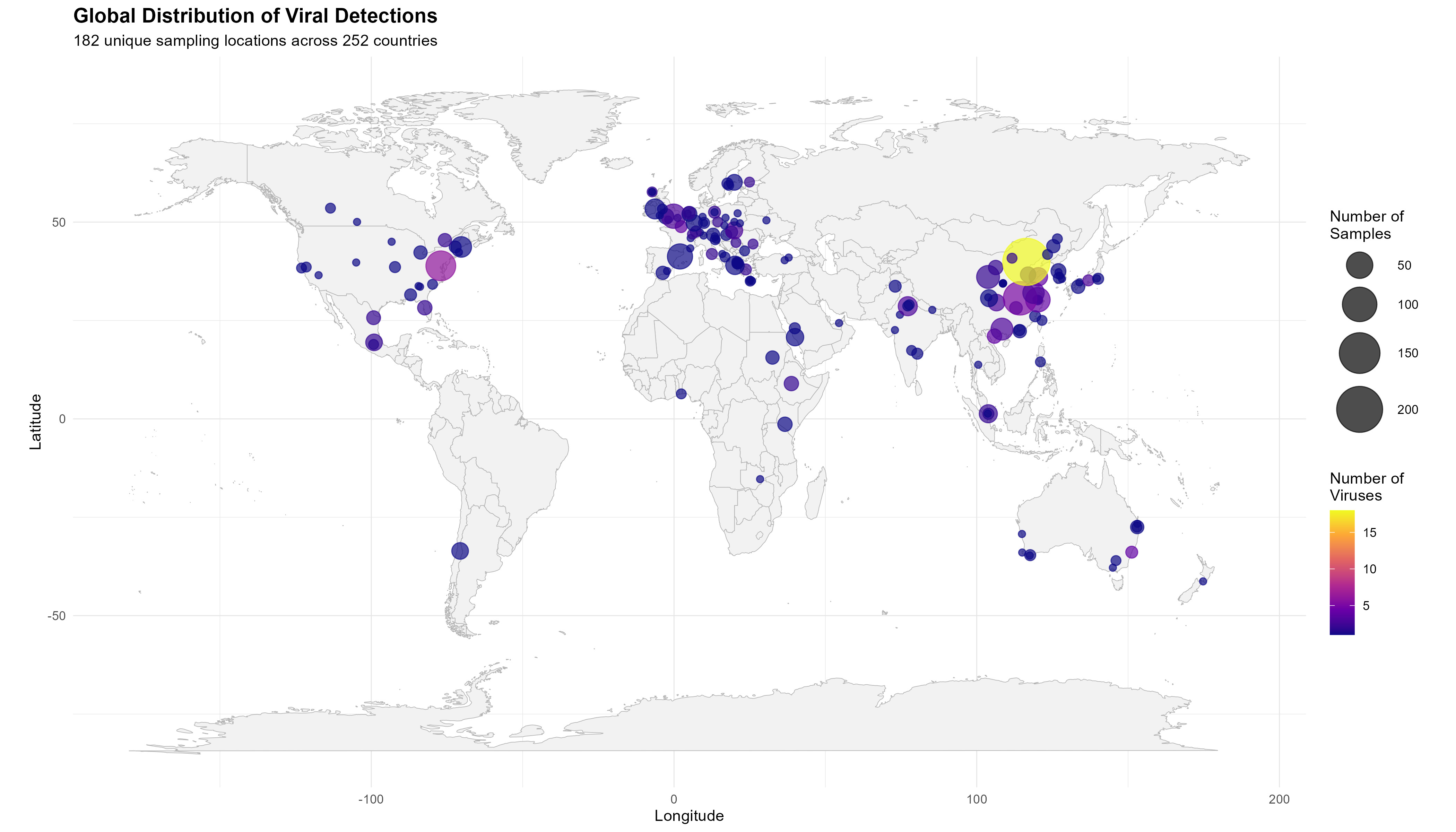
